# Cortical somatostatin interneuron subtypes form cell-type specific circuits

**DOI:** 10.1101/2022.09.29.510081

**Authors:** Sherry Jingjing Wu, Elaine Sevier, Giuseppe-Antonio Saldi, Sabrina Yu, Lydia Abbott, Da Hae Choi, Mia Sherer, Yanjie Qiu, Ashwini Shinde, Daniella Rizzo, Qing Xu, Irving Barrera, Vipin Kumar, Giovanni Marrero, Alvar Prönneke, Shuhan Huang, Bernardo Rudy, David A. Stafford, Evan Macosko, Fei Chen, Gord Fishell

**Affiliations:** Harvard Medical School, Blavatnik Institute, Department of Neurobiology, Boston, MA 02115, USA; Stanley Center for Psychiatric Research, Broad Institute of MIT and Harvard, Cambridge, MA 02142, USA; Department of Health Sciences, Bouvé College of Health Sciences, Northeastern University, Boston, MA 02115, USA; Department of Biology, Northeastern University, Boston, MA 02115, USA; Department of Behavioral Neuroscience, College of Science, Northeastern University, Boston, MA 02115, USA; Department of Biology, Brandeis University, Waltham, MA, USA; Center for Genomics & Systems Biology, New York University Abu Dhabi, Abu Dhabi, UAE; Neuroscience Institute, New York University School of Medicine, New York, NY, USA; Department of Molecular and Cell Biology, University of California, Berkeley, CA 94708

**Keywords:** Cortical interneurons, somatostatin, circuit, mouse genetics, optogenetics, rabies tracing, spatial transcriptomics

## Abstract

The cardinal interneuron classes are a useful simplification of cortical interneuron diversity, but such broad subgroupings glosses over the molecular, morphological, and circuit specificity of interneuron subtypes, most notably among the somatostatin interneuron class. The organizing principles by which the connectivity of these subtypes is specified are unknown. To address this knowledge gap, we designed a series of genetic strategies to target the breadth of somatostatin interneuron subtypes. Using these strategies to target three subtypes that span the entire cortical column, we examined their afferent and efferent connectivity. Our data demonstrated that each of these possesses remarkable reciprocal connectivity with the intracortical or corticofugal pyramidal classes, as well as parvalbumin interneurons. Even when two interneuron subtypes shared the same efferent target, their synaptic targeting proved selective for particular dendritic compartments. We thus provide evidence that subtypes of somatostatin cortical interneurons form cell-type specific cortical circuits.

## INTRODUCTION

The astonishing computational processing capacity of the mammalian cerebral cortex relies on the intricate connectivity between its two fundamental cell types, the glutamatergic excitatory neurons and GABAergic inhibitory interneurons. The layers of the cerebral cortex have long been recognized as being organized into a well-ordered circuitry comprised of distinct excitatory neuronal types across layers. It is less apparent whether cortical interneurons follow the same laminar organizational principles as the excitatory neurons, in part because of the absence of an obvious prescribed laminar distribution. Instead, diversity within cortical interneurons is primarily categorized by the expression of molecular markers and their targeting of subcellular compartments (Fishell and Kepecs, 2020; Tremblay et al., 2016). Thus, cortical interneurons can be coarsely grouped into four major cardinal classes expressing parvalbumin (*Pvalb*), somatostatin (*Sst)*, vasoactive intestinal peptide (*Vip*), or lysosome-associated membrane protein (*Lamp5*) genes (Yao et al., 2021). These four classes are generally attributed to serving non-overlapping circuit functions: feedforward inhibition, feedback inhibition, disinhibition, and “ bulk” or “tonic” inhibition (Kepecs and Fishell, 2014; Tremblay et al., 2016), each of which shows relatively stereotyped targeting of subcellular compartments, dendrites, somas or axons.

Here we focus on SST-expressing cortical interneurons that have previously been hypothesized to provide non-specific feedback inhibition to pyramidal neuron dendrites (Fino and Yuste, 2011; Fino et al., 2013; Karnani et al., 2014). Despite this, emerging evidence suggests that diversity within SST interneurons allows them to function in a more specific manner. Previous work examining the biophysical properties, morphology, and molecular markers has described at least three SST interneuron subtypes. The majority of SST interneurons are Martinotti cells, defined by an axonal plexus in L1, which can be further divided based on their morphology into fanning-out Martinotti cells with axons that ramify in both L2/3 and L1, and T-shaped Martinotti cells that ramify in L1 alone (Muñoz et al., 2017). In addition, there exists a population of non-Martinotti cells that target L4 instead of L1 (Ma et al., 2006; Muñoz et al., 2017; Nigro et al., 2018; Xu et al., 2013). Moreover, *in vivo* functional studies have shown that infragranular SST interneuron subtypes exert layer-specific control of sensory processing (Muñoz et al., 2017; Naka et al., 2019). Finally, recent advances in single-cell genomics have rapidly expanded our knowledge of interneuron transcriptomic diversity and unraveled additional SST interneuron subtypes (Mayer et al., 2018; Mi et al., 2018; Paul et al., 2017; Tasic et al., 2016, 2018; Yao et al., 2021). However, which properties best connote meaningful functional diversity is still a matter of debate. For example, in an effort to link transcriptionally-defined clusters (T) with historical classifications based on electrophysiology (E) and morphology (M), recent studies have utilized Patch-seq to collect and reconcile information on all three parameters from single neurons to define so-called MET types (Cadwell et al., 2016; Fuzik et al., 2016; Gouwens et al., 2020; Scala et al., 2021). Although this work made a direct and concerted effort to unify the various parameters that distinguish interneuron subtypes, it is unclear how the three features used for MET analysis relate to the functionality of the cell types that emerge from these classifications.

In this study, using SST interneurons as an exemplar, we propose connectivity as an organizing principle that synthesizes noisy cellular features into meaningful cell types. To test this hypothesis, we divided SST interneurons into eight transcriptomic subtypes (nine if one includes the CHODL type, representing SST-expressing long-range projecting neurons) and designed various genetic strategies for selectively targeting these different SST subtypes. Using a combination of spatial transcriptomics, morphological reconstructions of sparse-labeled neurons, single-molecule fluorescent *in situ* hybridization (smFISH), and electrophysiology, we validated that these subtypes represent the totality of SST interneurons in primary somatosensory and visual cortices. Furthermore, these characterizations revealed that each SST subtype possesses a unique laminar organization and stereotyped axonal projection pattern. To test whether the subtype-specific organization reflected discrete circuit motifs within local cortical networks, we used optogenetics to map the efferent connectivity of three major SST subtypes that are distributed in different layers onto local excitatory neurons. Intriguingly, each SST subtype had a distinct intralaminar or translaminar targeting pattern, as well as cell-type selective targeting of L5 pyramidal neurons and PV interneurons. Afferent mapping of two of the infragranular SST subtypes using monosynaptic rabies tracing provided further support that cell-type selective connectivity between different SST subtypes and excitatory neuron cell types is likely reciprocal. Finally, synaptic puncta analysis of two SST subtypes that innervate the same excitatory neuron cell type revealed that while they share a common efferent target, at the synaptic level, they likely gate distinct dendritic inputs. Our data demonstrate that SST interneurons can be divided into discrete subtypes that selectively contribute to cell-type specific circuits within the cortex. Taken together, this reveals an unanticipated precision of cortical interneurons in regulating the flow and excitability of cortical pyramidal cells.

## RESULTS

### SST interneurons subtypes are organized in layers

To assess the transcriptomic diversity of cortical SST interneurons, we took advantage of a single-nuclei RNA sequencing (snRNA-seq) dataset of cortical interneurons from mouse anterior lateral motor cortex (ALM) and primary visual cortex (V1) at postnatal day (P) 28 (Allaway et al., 2021). The use of snRNA-seq prevents stress-sensitive artifacts in gene expression and the selective loss of particular cell types compared to fluorescence-activated cell sorting (FACS) of whole cells (Bakken et al., 2018). This dataset utilized a Dlx5/6-Cre driver line, which enriches for all cortical interneurons and allows for the collection of the breadth of interneuron subtypes in accordance with their relative abundances. Interneurons derived from the medial and caudal ganglionic eminences (MGE and CGE, respectively) are clearly separated into distinct branches as visualized in a Uniform Manifold Approximation and Projection (UMAP) plot (Figure 1A, inset). Based on our data, SST interneurons can be divided into eight different subtypes (Github), and an additional CHODL subtype, which corresponds to nNos-expressing long-range projecting neurons (Figure 1B) (He et al., 2016; Paul et al., 2017; Tasic et al., 2016). Canonical correlation analysis (CCA) showed that this division closely matches with the supertypes of SST interneurons, as described in the recently published taxonomy of transcriptomic cell types of the isocortex and hippocampal formation (Yao et al., 2021) (analysis on Github). Therefore, with minor adjustments (e.g. combining two subtypes SST-Lpar1 and SST-Esm1 to SST-Nmbr see Github for details), we were able to adhere to the current nomenclature utilized by Allen Institute in classifying SST interneuron into eight subtypes (Figure 1A), each of which possesses distinct marker genes (Figure 1B).

**Figure 1.**
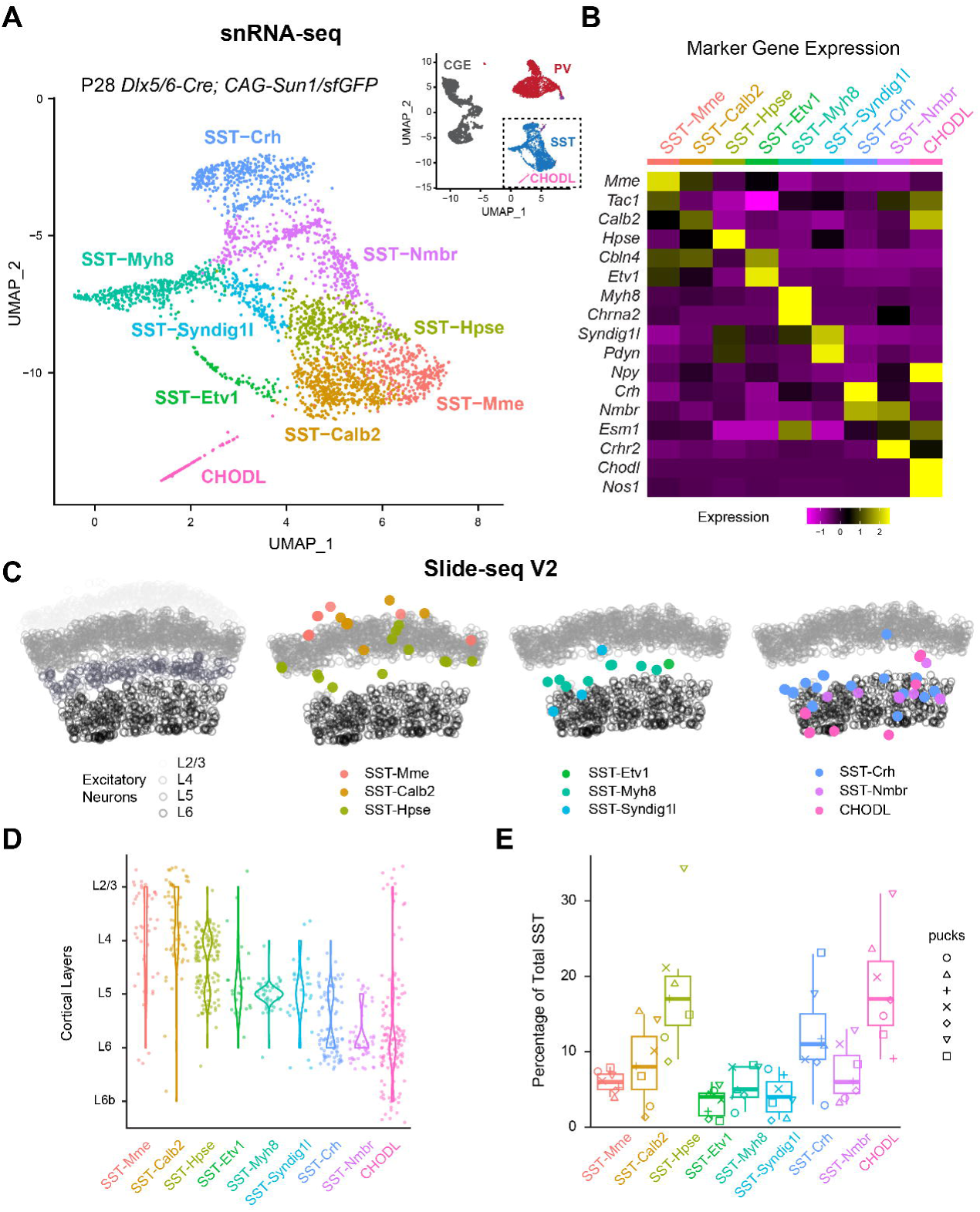
Spatial transcriptomic analysis reveals the laminar organization of eight SST interneuron subtypes. (A) UMAP visualization of P28 snRNA-seq of cortical interneurons (Allaway et al., 2021) illustrating the nine SST subtypes. Inset showing the UMAP of the entire dataset. CGE, caudal ganglionic eminence. (B) Heatmap showing the scaled expression of marker genes for each SST subtypes based on snRNA-seq data. (C) Robust cell type decomposition (RCTD) assignment of spatial clusters to different SST interneuron subtypes on a representative Slide-seq V2 experiment based on a scRNA-seq reference (see Methods). Gray circles represent the location of excitatory neurons in different layers for reference. (D) Violin plots demonstrate the laminar distribution of different SST subtypes identified in Slide-seqV2 experiments. (E) Boxplot showing the proportion of different SST subtypes out of total SST interneurons, found across seven Slide-seqV2 experiments. See also Figures S1-2, Tables S1-2.

To investigate the laminar distribution of these SST interneuron subtypes, we performed Slide-seq V2 experiments on the primary somatosensory cortex (S1) of ∼1-month-old mice (Stickels et al., 2021). With reference to scRNA-seq data (Yao et al., 2021), we used robust cell type decomposition (RCTD) (Cable et al., 2022) to detect the spatial distribution of different excitatory neuron cell types (Figure S1A,B) and the locations of each SST interneuron subtype (See Methods, Github). Interestingly, each SST subtype had a stereotyped laminar distribution: SST-Mme, SST-Calb2 are mainly found in upper layers; SST-Hpse reside in L4 and L5a; SST-Etv1, SST-Myh8, and SST-Syndig1l are all located in L5; and SST-Crh, SST-Nmbr, and CHODL are preferentially located within L6 (Figure 1C-D, S1C). To complement these observations, we performed single molecule fluorescent *in situ* hybridization (smFISH) against different marker genes for various SST subtypes in both S1 and V1 (Table S1). The laminar distribution of these marker genes confirms the Slide-seq V2 results, indicating that within these sensory cortical regions, specific SST subtypes reside in different cortical layers (Figure S2). These results also allowed us to estimate the relative proportion of different SST subtypes. In general, the results from Slide-seq V2 and smFISH agree well with each other. For instance, SST-Calb2 and SST-Crh subtypes were estimated to comprise ∼10% of the total SST interneuron population in S1 by both Slide-SeqV2 (Figure 1E) and smFISH (Figure S2). However, we did notice that Slide-seq V2 over-estimated the proportion of the CHODL subtype. In addition, the proportion of different SST subtypes in V1 can be estimated by the relative abundance in our snRNA-seq dataset (Table S2) or by smFISH (Figure S2) and are in concordance. Most of the SST subtypes are similar across these two sensory cortices, except that V1 contains a higher proportion of SST-Calb2 than S1 (∼19% in V1, ∼9% in S1) as estimated by smFISH (Figure S2). This correlates with the observation that the majority of the L4 SST interneurons in S1 belongs to SST-Hpse subtype (∼74% of SST-Hpse, ∼9% of SST-Calb2), while L4 of V1 is comprised by both SST-Calb2 (∼35%) and SST-Hpse (∼50%) (Figure S2), which has been previously shown by Patch-Seq of L4 SST interneurons in S1 and V1 respectively (Scala et al., 2019).

### Genetic targeting of different SST subtypes reveals stereotyped axonal projection patterns

Based on marker gene expression, we designed direct and intersectional genetic strategies to target either one or a combination of multiple SST interneuron subtypes (Table S3). These genetic strategies revealed the distinct laminar organization and axonal projection patterns of different SST subtypes that were largely consistent between S1 and V1 (Figure 2A, S3A). One exception to this trend is the SST-Calb2 subtype. The intersectional strategy of *Calb2^Cre^;Sst^FlpO^*primarily targets SST-Calb2 subtype in L2/3 and L5a, but not in L4 of S1 (Figure 2A). By comparison, the same genetic strategy showed that the SST-Calb2 subtype in V1 is distributed throughout L2/3 to L5a (Figure S3A), which is consistent with the smFISH results (Figure S2) and a previous publication (Scala et al., 2019). Instead, L4 of S1 is primarily populated by the SST-Hpse subtype, which after the age of P10 (due to germline expression of *Hpse* gene and the postnatal onset of *Hpse* expression in SST interneurons) can only be targeted by injection of recombinant adeno-associated viruses (AAVs), which drives Cre recombinase-dependent (Cre-ON) expression of reporter protein under a Dlx enhancer (Dimidschstein et al., 2016). SST-Hpse interneurons have extensive axons that arborize within L4, which results in the striking labeling of barrel fields in S1. Single-cell reconstruction confirms that the axons of SST-Hpse primarily target L4 in both S1 and V1, often with a collateral to L1. In S1, the axon of one SST-Hpse interneuron can fill an entire barrel field (Figure 2B, S3B). Therefore, the SST-Hpse subtype is a L4-targeting non-Martinotti cell that resides in L4 and L5a of both sensory cortices (Gouwens et al., 2020; Muñoz et al., 2017; Naka et al., 2019; Scala et al., 2019; Xu et al., 2013). As an alternative to targeting with a viral strategy, the *Pdyn^CreER^;Npy^FlpO^* Cre-ON/Flp-ON intersectional strategy can also be used to target SST-Hpse subtype, although this strategy in addition labels a subset of SST-Calb2. In addition, by crossing this compound allele with a Cre-ON/Flp-OFF reporter line, one can selectively target the SST-Syndig1l subtype, whose morphology corresponds to L5a T-shaped Martinotti cells (Figure 2, S3) (Muñoz et al., 2017; Nigro et al., 2018). Another SST subtype that resides in L5a is the SST-Etv1 subtype, which resembles a fanning-out Martinotti shape and can be partially targeted using an *Etv1^CreER^; Sst^FlpO^* intersectional strategy. Within L5b and 6, the *Chrna2-Cre* allele can be used to target the SST-Myh8 subtype, which also exhibits a T-Shaped Martinotti morphology (Figure 2, S3), as previously described (Hilscher et al., 2017).

**Figure 2.**
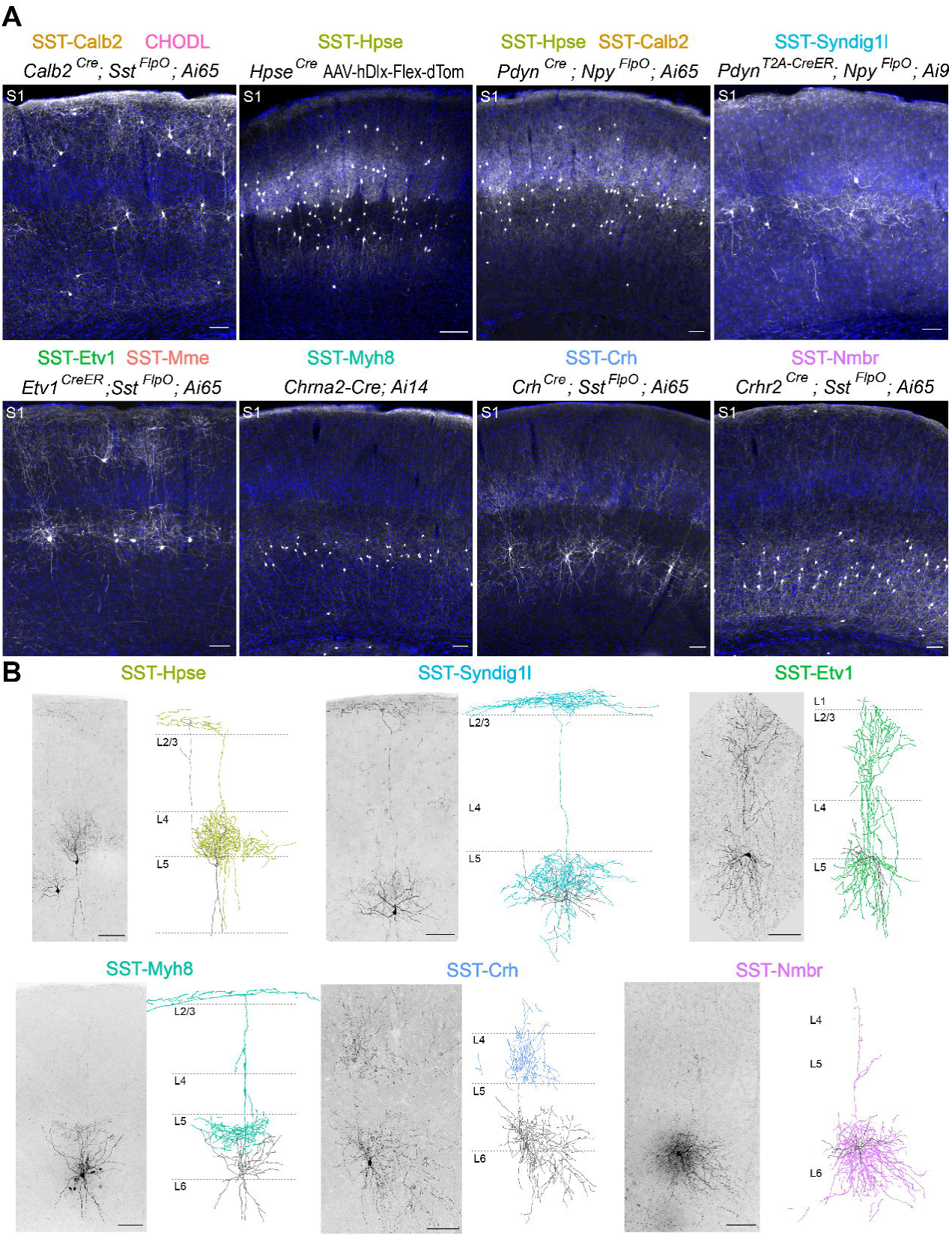
Genetically targeted SST subtypes showed stereotypical laminar distribution and morphology. (A) Representative images of genetic strategies for targeting SST subtypes in S1, contrasted with DAPI staining to allow visualization of their laminar distribution. All images were taken from 1-3 month old mice. *Ai9* reporter line is used here as a Cre-ON/Flp-OFF strategy due to the FRT sites flanking the LoxP cassette are retained in this mouse line (Madisen et al., 2010). With this reporter line, SST-Hpse interneurons were occasionally observed in *Pdyn^T2A-CreER^; Npy^FlpO^; Ai9* strategy, likely due to incomplete FlpO recombination, though not noted in this representative image. All panels except that showing SST-Hpse utilized entirely genetic strategies for targeting different SST subtypes. For SST-Hpse labeling, rAAV9-hDlx-Flex-dTomato virus was stereotaxically injected in *Hpse^Cre^* mice in S1 at 1-month-old and examined 13 days post-injection. Note that *Etv1^CreER^;Sst^FlpO^*intersectional strategy may target some SST-Calb2 interneurons as well. Scale bars, 100 µm. (B) Sparse labeling and Neurolucida reconstructions of selective SST subtypes in S1. Images of genetic labeled or biocytin-filled SST interneuron is shown to the left of the Neurolucida reconstruction of single-neuron morphology. SST-Etv1 interneuron is labeled by *Etv1^CreER^; Sst^FlpO^; RC::FPSit* genetic strategy. SST-Hpse and SST-Syndig1l interneurons are both labeled by *Pdyn^T2A-CreER^; Ai14* strategy and differentiated by their unique morphology. SST-Crh interneuron is labeled by *Crh^Cre^; Sst^FlpO^; RC::FPSit*. SST-Myh8 and SST-Nmbr are both labeled by biocytin-filling. All reconstructions were derived using sparse labeling of specific SST subtypes from P25-73 mice. Scale bars, 100 µm. See also Figures S3-5, Tables S3-6.

Little has been previously reported about SST interneuron diversity in L6. We identified two strategies for targeting two L6 SST subtypes, each of which has distinct features. The *Crh^Cre^;Sst^FlpO^*intersectional strategy targets the SST-Crh subtype, which are L4-targeting non-Martinotti cells that reside within L5b and L6 (Naka et al., 2019). The *Crhr2^Cre^;Sst^FlpO^*intersectional strategy targets the SST-Nmbr subtype that resides almost exclusively in L6. The axons of these cells remain primarily in deep layers despite occasionally extending thin collateral towards L1, suggesting that they are also non-Martinotti cells (Figure 2, S3). This morphology was also observed in previously published single-cell reconstructions of L6 SST interneurons (Gouwens et al., 2020). For SST subtypes not highlighted here, we have included a list of genetic targeting strategies (Table S4) and a summary of our current understanding of the putative SST subtypes targeted using each genetic approach (Table S4). Additional images of sparse labeling of each targeted SST subtype can be found at https://doi.org/10.7910/DVN/NQDIPG. Based on the transcriptomic clusters (Table S4), our sparse labeling in general agrees with the published single-neuron reconstructions of the corresponding transcriptomic clusters (Gouwens et al., 2020).

To assess the completeness and coverage of these genetic strategies, we performed smFISH against *Sst* mRNA for quantification of genetic labeling in S1 and V1 (Figure S4, S5). In general, most genetic strategies label the expected proportion of SST subtypes. For example, the *Calb2^Cre^;Sst^FlpO^* strategy labels ∼22% of total SST interneurons in V1 (Figure S4), which is consistent with the prevalence of the SST-Calb2 subtype as estimated by snRNA-seq (∼20% of SST interneurons in V1, Table S2) and smFISH (∼19%, Figure S2). In addition, consistent with that mentioned above, the *Calb2^Cre^;Sst^FlpO^* strategy labels ∼8% more SST interneurons in V1 compared to S1, in accordance with their greater abundance in the former. Similarly, the *Crhr2^Cre^;Sst^FlpO^*strategy labels ∼11% of total SST interneurons in V1 (Figure S4) and thus correlates well with the size of the SST-Nmbr population, which by snRNA-seq is estimated to be ∼13% of total SST interneurons in this area (Table S2). However, when the *Crh^Cre^; Sst^FlpO^* genetic targeting strategy is used, the SST-Crh subtype seems to be underrepresented, as this only results in the labeling of ∼2-3% of total SST interneurons in both S1 and V1 (Figure S4), while the SST-Crh subtype is estimated to comprise >10% of total SST interneurons by Slide-seq V2 (Figure 1), snRNA-seq (Table S2) and smFISH (Figure S2).

To evaluate the specificity of four of the genetic strategies, we performed smFISH experiments using selected marker genes. Overall, each genetic strategy showed the expected expression pattern of the selected marker genes in each of the targeted SST subtypes (Figure S5). For example, both *Chrna2-Cre* and *Crhr2^Cre^;Sst^FlpO^* labeled SST interneurons with low levels of *Calb2* and *Hpse* transcripts (Figure S5). However, the majority of the marker genes are not binary classifiers for each SST subtype. Instead, they showed graded expression across the examined SST subtypes (Figure 1B, Table S1). Notably, due to the high sensitivity of the smFISH method, a low level of transcripts is often detected. For example, *Pdyn* gene is expressed at a low level in the SST-Calb2 subtype (Table S1), resulting in a high percentage of *Pdyn*+ *Calb2^Cre^;Sst^FlpO^*labeled SST interneurons (Figure S5A). Therefore, to undertake a thorough characterization of specificity and coverage of each of the genetic strategies utilized, would require a quantitative analysis involving the smFISH multiplexing of upwards ∼20 genes. A further caveat associated with strategies involving tamoxifen-dependent labeling is that different proportions of SST subtypes are labeled in specific experiments, depending on the recombination efficiency, as a result of the graded expression within the targeted populations. For example, immunostaining against Calretinin (the protein product of the *Calb2* gene) suggested that the *Etv1^CreER^; Sst^FlpO^* intersectional strategy, in addition to targeting SST-Etv1 populations, labels the SST-Mme subtype (that expresses a low-level of *Calb2*) (Figure S5B), due to the graded expression of *Etv1* gene in different SST subtypes.

A notable outcome of our genetic analysis was that it revealed that each SST transcriptomic subtype has a stereotypical laminar location and an associated axonal projection pattern (see also (Gouwens et al., 2020). We likewise wondered whether they exhibited stereotyped electrophysiological properties. To test this, we decided to focus on three major SST interneuron subtypes that tiled the cortical column: the SST-Calb2, the SST-Myh8 and the SST-Nmbr subtypes, targeted with the *Calb2^Cre^;Sst^FlpO^*, *Chrna2-Cre* and *Crhr2^Cre^;Sst^FlpO^* alleles, respectively. These three SST subtypes each reside in different cortical layers and have distinct morphologies. SST-Calb2 interneurons are fanning-out Martinotti cells found in L2/3 to L5, SST-Myh8 interneurons are T-shaped Martinotti cells concentrated in L5b, and SST-Nmbr interneurons are L6 non-Martinotti cells whose axons primarily arborize extensively in deep layers (Figure 2) (Gouwens et al., 2020; Hilscher et al., 2017). Previous studies have characterized three major electrophysiological profiles for SST interneurons: adapting regular spiking, quasi-fast spiking, and low-threshold spiking (LTS) (He et al., 2016; Ma et al., 2006; Naka et al., 2019; Nigro et al., 2018; Xu et al., 2013). These electrophysiological types correlate with previously described SST subtypes in several transgenic lines (GIN, X94, and X98, respectively), as well as morphological parameters (Ma et al., 2006; Nigro et al., 2018; Xu et al., 2013), but recent efforts to link transcriptomic clusters with electrophysiology have found significant variability across transcriptional types (Gouwens et al., 2020; Scala et al., 2019). To address whether these SST subtypes have particular biophysical identities, we analyzed 11 electrophysiological parameters from genetically labeled SST-Calb2, SST-Myh8, and SST-Nmbr interneurons (Table S5). To ensure our results are comparable with previous studies, we restricted our analysis to S1. Overall, all three subtypes displayed regular-spiking firing patterns with adaption (Figure S3C). While SST-Myh8 interneurons displayed rebound burst firing, they did not have the characteristic high input resistance, low action potential (AP) threshold, or high adaptation index of low threshold spiking (LTS) cells. We trained a k-nearest neighbor classifier on our dataset and found that SST-Calb2 and SST-Nmbr interneurons were predicted with >80% accuracy, but SST-Myh8 interneurons were mixed with SST-Calb2, likely due to rebound firing in some SST-Calb2 cells (Figure S3B,C). SST-Nmbr interneurons were primarily distinguished by their higher firing frequency (Figure S3C). Notably, SST-Calb2 interneurons are found in both L2/3 and L5a in S1. To address whether SST-Calb2 interneurons are a continuum of one cell type or two distinct cell types in different layers, we compared the intrinsic electrophysiological properties of SST-Calb2 interneurons in these two layers. We found that across most parameters, SST-Calb2 interneurons in L2/3 were indistinguishable from those in L5a, with the exception that L5a SST-Calb2 interneurons were slightly more adapting (Table S6). In summary, despite their distinct electrophysiological properties, the three SST subtypes are better discerned by other features such as laminar location, morphology and axonal projection patterns.

### Laminar positioning of SST subtypes partially predicts their output connectivity

Given that SST-Calb2, SST-Myh8, and SST-Nmbr are organized in distinct cortical layers, we hypothesized that they may form laminar-specific circuits. To test this, we used our genetic strategies (*Sst^Cre^;Sst^FlpO^*for pan-SST cells, *Calb2^Cre^;Sst^FlpO^* for SST-Calb2, *Chrna2-Cre;Sst^FlpO^* for SST-Myh8, and *Crhr2^Cre^;Sst^FlpO^*for SST-Nmbr) in the context of the intersectional reporter line *Ai80*, which allows for recombinase-mediated expression of a channelrhodopsin variant, CatCh, in all SST interneurons or a particular SST subtype, accordingly (Figure 3A) (Daigle et al., 2018). Using this approach, we performed optogenetic-assisted circuit mapping experiments in V1.

**Figure 3.**
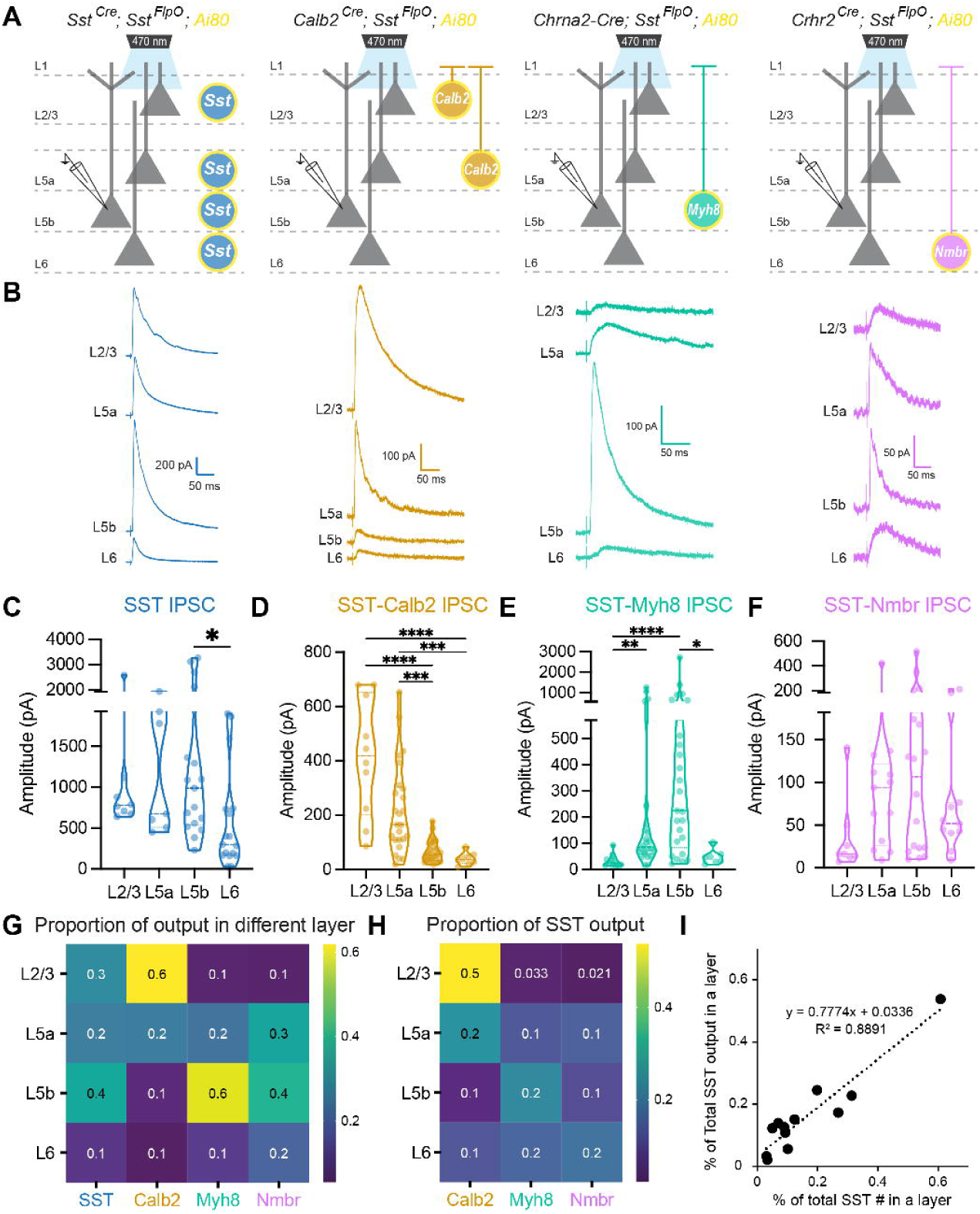
Laminar positioning correlates with SST subtype innervation. (A) Recording scheme. Pan-SST interneurons or three SST subtypes, SST-Calb2, SST-Myh8, SST-Nmbr, were genetically targeted to express CatCh by crossing with the *Ai80* reporter line. Postsynaptic IPSCs were recorded from pyramidal neurons across layers in response to 1 ms light stimulation. (B) Example average traces from pyramidal neurons across layers in response to pan-SST stimulation (left) and individual SST subtypes (three right panels). (C) Violin plot quantifying the evoked IPSC amplitude upon pan-SST interneuron stimulation. Y-axis line break represents the largest 75^th^ quartile value, here from L5a at 1923 pA, with the bottom section representing 75% of the distribution. Output to L5b excitatory neurons were significantly stronger than L6 (p = .0277), but all other comparisons were not significant by Kruskal-Wallis test with Dunn’s correction. (D) Amplitude of evoked IPSC from SST-Calb2 interneurons. IPSC in L2/3 and L5a was significantly higher than in L5b and L6 (L2/3 vs. L5b and L2/3 vs. L6 p <.0001, L5a vs. L5b p = .0007, L5a vs. L6 p = .0009). IPSC in L2/3 was not significantly different from L5a (p = .505). (E) Amplitude of evoked IPSC from SST-Myh8 interneurons. IPSC in L5b was not significantly higher than L5a (p = .8067), but was significantly higher than L2/3 (p < .0001) and L6 (p = .0161). IPSC in L5a was significantly higher than in L2/3 (p = .0013) but not L6 (p = .2778). (F) Amplitude of evoked IPSC from SST-Nmbr interneurons. All comparisons not significant (L2/3 vs L5a p = .2317, L2/3 vs. L5b p = .1147, L2/3 vs. L6 p = .6637. L5a vs. L5b, L5a vs. L6, L5b vs. L6 all p > .9999). (G) Heatmap of ratio of median evoke IPSC amplitude for pan-SST interneurons or for individual SST subtypes across layers. Data were normalized across columns, where the value represents the ratio between the median evoked IPSC amplitude in a particular layer compared to the total median inhibitory output of that SST type across layers (i.e. 60% of inhibitory output of SST-Calb2 cells was in L2/3, 20% in L5a etc.) (H) Heatmap of the proportion of inhibition from individual SST subtype as compared to the inhibition from pan-SST interneurons in different layers. (I) Plot showing that percentage of individual SST subtype out of total number of SST interneuron found in a particular layer (x axis) is correlated with the proportion of the inhibitory output by individual SST subtype out of pan-SST interneurons in that layer (y axis). See also Figure S6.

We first examined the output of SST interneurons as a general class, and found that SST interneurons strongly inhibit all layers, with the smallest response in L6 (Figure 3B-C). Note that the level of inhibition does not correlate with the abundance of SST interneurons found in each layer (Figure S4, S6E). Compared with the other layers, L6 seems to receive disproportionately less inhibition from SST interneurons (Figure S6E), suggesting that L6 excitatory neurons either receive less innervation or form weaker synapses with SST interneurons compared to excitatory neurons in other layers (Campagnola et al., 2022). We then tested whether individual SST subtypes also selectively target specific cortical layers. We found that corresponding with their laminar positioning, SST-Calb2 interneurons primarily innervate L2/3 and L5a (Figure 3B,D). SST-Myh8 interneurons likewise innervated their resident layer L5b, as well as pyramidal cells within the adjacent L5a layer (Figure 3B,E). Surprisingly, SST-Nmbr interneurons did not show preferential laminar targeting despite their cell bodies being mostly restricted to L6 (Figure 3B,F). The median evoked IPCS amplitude from SST-Nmbr subtype is larger in L5, as compared to L6 (Figure 3G), consistent with L6 excitatory cells receiving less SST-mediated inhibition than other layers. As a general trend, when we examined the contribution from each SST subtype in proportion to the pan-SST output, as measured in the soma. each subtype consistently contributes most to the overall inhibition of its resident layer (Figure 3H, Figure S6A-D). In fact, the percentage of each SST subtype found in each layer correlates well with the portion of their contribution to the total inhibitory output by SST interneurons in that layer (Figure 3I). However, the strength of inhibition was not distributed equally across each SST subtype. SST-Calb2 subtype tends to form stronger inhibition as compared to the other two subtypes (Figure S6F). This could be due to differences in the strength of the inhibitory synapses formed, the number of synapses per cell, or the input resistance of the postsynaptic neuron.

Notably, compared to evoked IPSCs from pan-SST stimulation, all three subtypes evoked significantly lower responses (Figure S6A-D). To assess the combined contribution of these three SST subtypes to the pan-SST response, we compared simulated linear combinations of each subtype with the median pan-SST response for a particular cortical layer using a hierarchical bootstrapping method (see Methods). We found that the combined evoked IPSC amplitude from SST-Calb2, SST-Myh8, and SST-Nmbr was smaller than the pan-SST response in 75-85% of the trials across all layers (Figure S6E). To estimate the proportion of the pan-SST response that could be attributed to the output of these three subtypes, we took the ratio of a linear combination of each subtype compared to a pan-SST evoked amplitude (Figure S6F). The highest proportion was in L2/3 pyramidal neurons with a median ratio of 65.31%, followed by L6 at 59.54%, L5a at 59.51%, and L5b at 55.96%. This is not surprising given that the combination of these three SST subtypes constitutes ∼46% of all SST interneurons, and depending upon the layer, varies from 40-60% in their relative abundance (Figure S6H).

Taken together, these results suggest that in aggregate each SST subtype contributes most to its resident layer, although some subtypes may target cells in other layers more strongly. It is particularly intriguing that Martinotti cells such as SST-Calb2 and SST-Myh8, despite their axons being largely restricted to L1, can still selectively target excitatory neurons in their resident layer. This suggests that there is a mechanism for SST interneurons to recognize the dendrites of pyramidal cells whose soma they are proximal to.

### SST subtypes selectively target IT and PT neurons within L5

All three SST subtypes examined innervate L5, which contains two major types of pyramidal neurons: intratelencephalic (IT) neurons that project within the cortex and striatum; and pyramidal tract (PT) neurons that extend their axon subcerebrally to several targets including the tectum, brainstem, and spinal cord. We therefore wondered whether these SST subtypes selectively innervate specific pyramidal neuron types within the same layer. To address this, we injected rAAV2-retro-hSyn-mScarlet into either the ipsilateral retrosplenial cortex (Rs) or superior colliculus (SC) to retrogradely label IT and PT neurons in V1, respectively (Figure 4A). We then recorded optogenetically evoked IPSCs from virally labeled IT and PT neurons in L5 using the same genetic strategy for targeting SST subtypes (Figure 4B).

**Figure 4.**
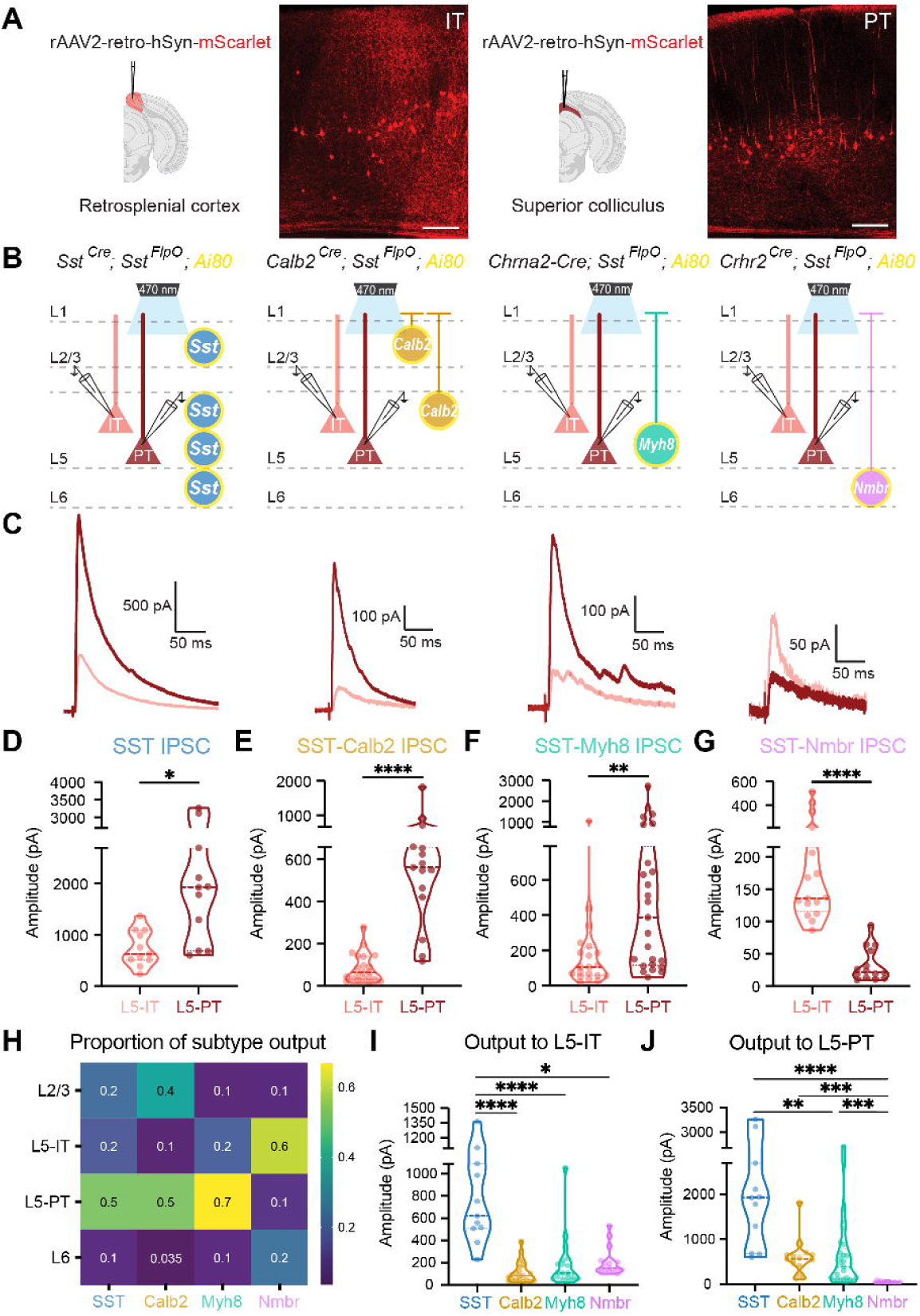
SST subtypes differentially target IT vs. PT pyramidal neurons in L5. (A) Strategy for targeting IT and PT pyramidal neurons by injecting rAAV2-retro-hSyn-mScarlet into the retrosplenial cortex (Rs) or superior colliculus (SC), respectively. Representative images of mScarlet-labeled PT and IT neurons. Scale bars, 100 µm. (B) Recording scheme. Pan-SST interneurons or three SST subtypes, SST-Calb2, SST-Myh8, SST-Nmbr, were genetically targeted to express CatCh by crossing with the *Ai80* reporter line. Postsynaptic IPSCs are recorded from IT or PT neurons in response to 1 ms light stimulation. (C) Representative average traces of evoked IPSC in IT (pink) and PT (red) neurons upon stimulation of pan-SST, SST-Calb2, SST-Myh8, and SST-Nmbr interneurons. (D) Violin plot of evoked IPSC amplitude upon optogenetic stimulation of pan-SST interneurons in L5-IT and L5-PT neurons. IPSC amplitude in L5-PT neurons was significantly greater than in L5-IT neurons (Kolmogorov-Smirnov, p = .0233). (E) As in (D) for SST-Calb2 interneurons. IPSC amplitude was significantly greater in L5-PT neurons than in L5-IT neurons (Mann-Whitney, p < .0001). (F) As in (D) for SST-Myh8 interneurons. IPSC amplitude in L5-PT neurons was significantly greater than in L5-IT neurons (Mann-Whitney, p = .0027). (G) As in (D) for SST-Nmbr interneurons. IPSC amplitude in L5-IT neurons was significantly greater than in L5-PT neurons (Mann-Whitney, p < .0001). (H) Heatmap of the proportion of inhibition from individual SST subtype as compared to the inhibition from pan-SST interneurons in different layers and pyramidal neuron cell types. (I) Evoked IPSC in L5-IT pyramidal neurons. Pan-SST interneuron response was greater than all subtypes (KruskalLJWallis test with Dunn’s correction for multiple comparisons, pan-SST vs. SST-Calb2 and SST-Myh8 p < .0001, vs. SST-Nmbr p = .03). (J) Violin plot of evoked IPSC in L5-PT pyramidal neurons. Pan-SST interneuron response was not significantly greater than SST-Calb2 (p = .1), but was greater than SST-Myh8 (p = .008) and SST-Nmbr (p < .0001). SST-Calb2 and SST-Myh8 responses were still significantly greater than SST-Nmbr (p = .0001, .0003 respectively). KruskalLJWallis test with Dunn’s correction for multiple comparisons. See also Figure S6.

We first tested the efferent connectivity of pan-SST interneurons to IT and PT neurons and found that SST interneurons strongly inhibit both types but have stronger output onto PT neurons (Figure 4C-D). Upon examination of individual SST subtypes, each one showed a clear innervation bias towards specific L5 pyramidal neuron cell types. Both SST-Calb2 and SST-Myh8 interneurons preferentially target L5-PT neurons, while SST-Nmbr interneurons preferentially inhibit L5-IT neurons (Figure 4C,E-G).

Compared across layers, this time with L5 separated into IT and PT neurons, pan-SST interneurons inhibited all layers and both cell types tested, although as noted above, significantly weaker in L6 (Figure 4H, Figure S6I). By comparison, clear subtype-specific patterns emerged for individual SST subtypes. SST-Calb2 strongly targeted L2/3 and L5-PT neurons, SST-Myh8 primarily targeted L5-PT neurons, and SST-Nmbr preferentially targeted L5-IT neurons (Figure 4H, Figure S6I-L). Note that SST-Calb2 and SST-Myh8 outputs to L5-PT neurons were not significantly different, showing that these two subtypes innervate L5-PT neurons with similar strength at a population level, while the inhibition from SST-Nmbr to L5-PT neurons was negligible (Figure 4I, Figure S6M-N). With regard to L5-IT neurons, outputs from these three SST subtypes are all relatively weak, although, amongst the three, SST-Nmbr was still the strongest (Figure 4J, Figure S6M-N).

To predict the contribution of these three types to the pan-SST inhibition, we repeated the hierarchical bootstrapping and linear combination simulations described above. We found that the combined IPSC simulated responses were 74.84% less in L5-PT cells and 77.68% less in L5-IT cells than observed upon Pan-SST stimulation (Figure S6O). To estimate the proportion of the pan-SST inhibition that could be attributed to SST-Calb2, SST-Myh8, and SST-Nmbr outputs, we took the ratio of a linear combination of each subtype compared to a pan-SST evoked amplitude (Figure S6P). For L5-PT neurons the median contribution was 61.48%, and for L5-IT neurons the median contribution was 66.23%, indicating that these subtypes account for approximately two-thirds of the total SST inputs to both L5-PT and L5-IT pyramidal neurons. However, in both bootstrapping analyses our assumption that these inputs are linearly summated needs to be further investigated.

### SST subtypes differentially inhibit PV interneurons across layers

As we observed a high degree of specificity between three SST subtypes and excitatory neurons, we wondered whether they formed specific connections with inhibitory neurons as well. Previous studies suggest that except for themselves, SST interneurons broadly inhibit all other cardinal classes of interneurons (Pfeffer et al., 2013). There is already evidence that non-Martinotti SST interneurons innervate L4 PV interneurons more strongly than other SST interneurons (Xu et al., 2013). To test for selective outputs to PV interneurons from these three SST subtypes, we injected an AAV expressing GFP under the control of a PV-specific enhancer (Vormstein-Schneider et al., 2020) into the various SST subtype-specific driver lines crossed with *Ai80.* We then proceeded to record from virally labeled PV interneurons (Figure 5A-B). We found that SST-Calb2 interneurons strongly targeted PV interneurons in the superficial layers, while SST-Myh8 and SST-Nmbr interneurons targeted PV interneurons weakly in infragranular layers but not at all in the superficial layers (Figure 5C-E). To compare this to previous studies demonstrating the innervation of L4 PV interneurons by non-Martinotti cells, we also tested the output of the SST interneurons labeled using the *Pdyn^Cre^;Npy^FlpO^*strategy. This strategy primarily targets the L4-targeting non-Martinotti SST-Hpse subtype, as well as some SST-Calb2 cells. As expected, SST interneurons targeted using this genetic strategy strongly innervated L2/3 and L4 PV interneurons (data not shown).

**Figure 5.**
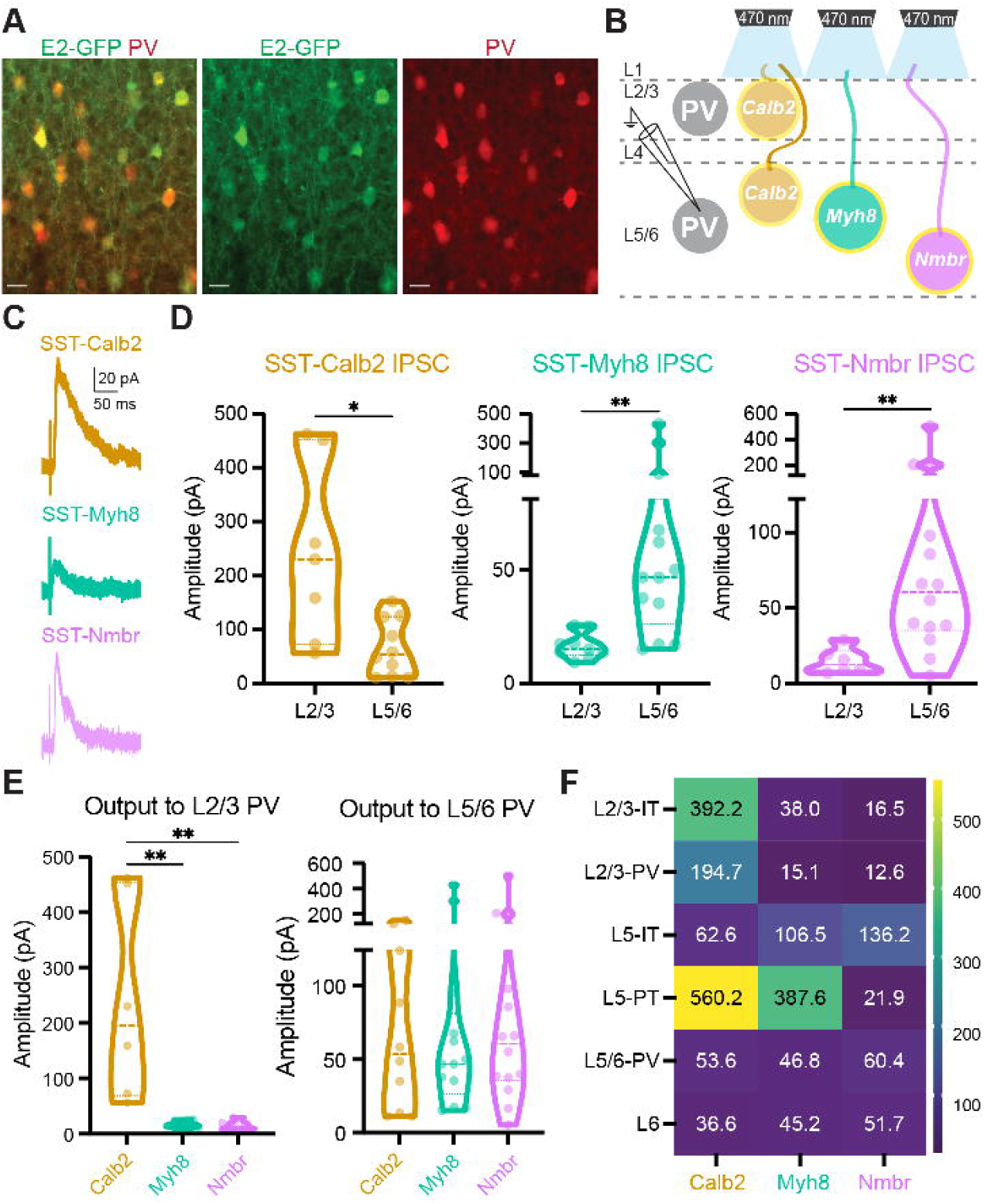
SST subtypes differentially innervate PV interneurons. (A) Representative image of E2-GFP injection in V1 labeling PV interneurons. Scale bars, 5 µm. (B) Recording scheme. Three SST subtypes, SST-Calb2, SST-Myh8, SST-Nmbr, were genetically targeted to express CatCh by crossing with the *Ai80* reporter line. IPSCs were recorded from PV neurons in response to 1 ms light stimulation. (C) Representative traces of IPSCs in PV interneurons in response to optogenetic stimulation of SST-Calb2, SST-Myh8, or SST-Nmbr interneurons. (D) Quantification of IPSC amplitudes upon optogenetic stimulation. IPSC from SST-Calb2 interneurons was significantly greater in L2/3 PV than in L5/6 PV interneurons (p = .0140), while IPSC from SST-Myh8 and SST-Nmbr was greater in L5/6 PV interneurons (p = .0037, p = .0014). Kolmogorov-Smirnov test. (E) IPSCs from SST-Calb2, SST-Myh8, and SST-Nmbr interneurons in L2/3 (left) and L5/6 PV interneurons (right). IPSCs from SST-Calb2 interneurons to L2/3 PV was significantly greater than from SST-Myh8 (p = .0071) and SST-Nmbr (p = .0047). Outputs to L5/6 PV interneurons were not significantly different across SST subtypes. Kruskal-Wallis test. (F) Heatmap of median evoked IPSC amplitude (pA) from each SST subtype across pyramidal neurons and PV interneurons in different layers.

A heatmap of median evoked IPSC amplitude summarizes the selective output patterns of the three SST subtypes across different layers and cell types (Figure 5F). The three SST subtypes we studied proved to have a combination of shared and distinct targets. SST-Calb2 interneurons targeted L2/3 and L5-PT pyramidal neurons and L2/3 PV interneurons and SST-Myh8 interneurons targeted L5-PT pyramidal neurons. While SST-Nmbr interneurons targeted L5-IT neurons specifically, none of the three subtypes provided strong input to L5-IT or L6 pyramidal neurons compared to the pan-SST response.

### Two infragranular SST subtypes receive reciprocal selective excitatory neuron inputs

As our optogenetic experiments demonstrated that different SST subtypes had selective output connectivity, we wondered whether they also received selective input connectivity. To test this, we performed monosynaptic rabies tracing on two closely positioned infragranular SST subtypes, SST-Myh8 and SST-Nmbr. To restrict starter cells to specific SST subtypes, we utilized AAV-helper viruses that allow Cre-dependent expression of TVA receptor for the infection of EnvA-pseudotyped rabies virus (RV), and G protein for the replication and monosynaptic transport of RV. Specifically, for targeting SST-Myh8 interneurons, we injected AAV-helper viruses (AAV-hSyn-DIO-TVA-GFP-N2cG) (Pouchelon et al., 2021) in *Chrna2-Cre* mice at an early developmental age (P2-5), due to the downregulation of *Chrna2-Cre* expression around the third postnatal week. We subsequently injected N2c-RV-mCherry virus at P22-42 for S1 and P56-79 for V1 (Figure 6A). For targeting SST-Nmbr interneurons, we co-injected AAV-helper viruses (AAV-Dlx-DIO-TVA and AAV-Dlx-DIO-GFP-N2cG) with N2c-RV-mCherry in 1-3 month old *Crhr2^Cre^*mice within both S1 and V1 (Figure 6A). The use of the mDlx5/6 enhancer (Dimidschstein et al., 2016) in the AAV-helper viruses is necessary for selective targeting SST-Nmbr interneurons because a subset of L2/3 pyramidal neurons also express *Crhr2* gene (data not shown). Retrogradely traced presynaptic neurons were examined 10-14 days after RV injection. As expected, GFP-positive starter cells were mainly found in L5b for SST-Myh8 and L6 for SST-Nmbr (Figure S7A).

**Figure 6.**
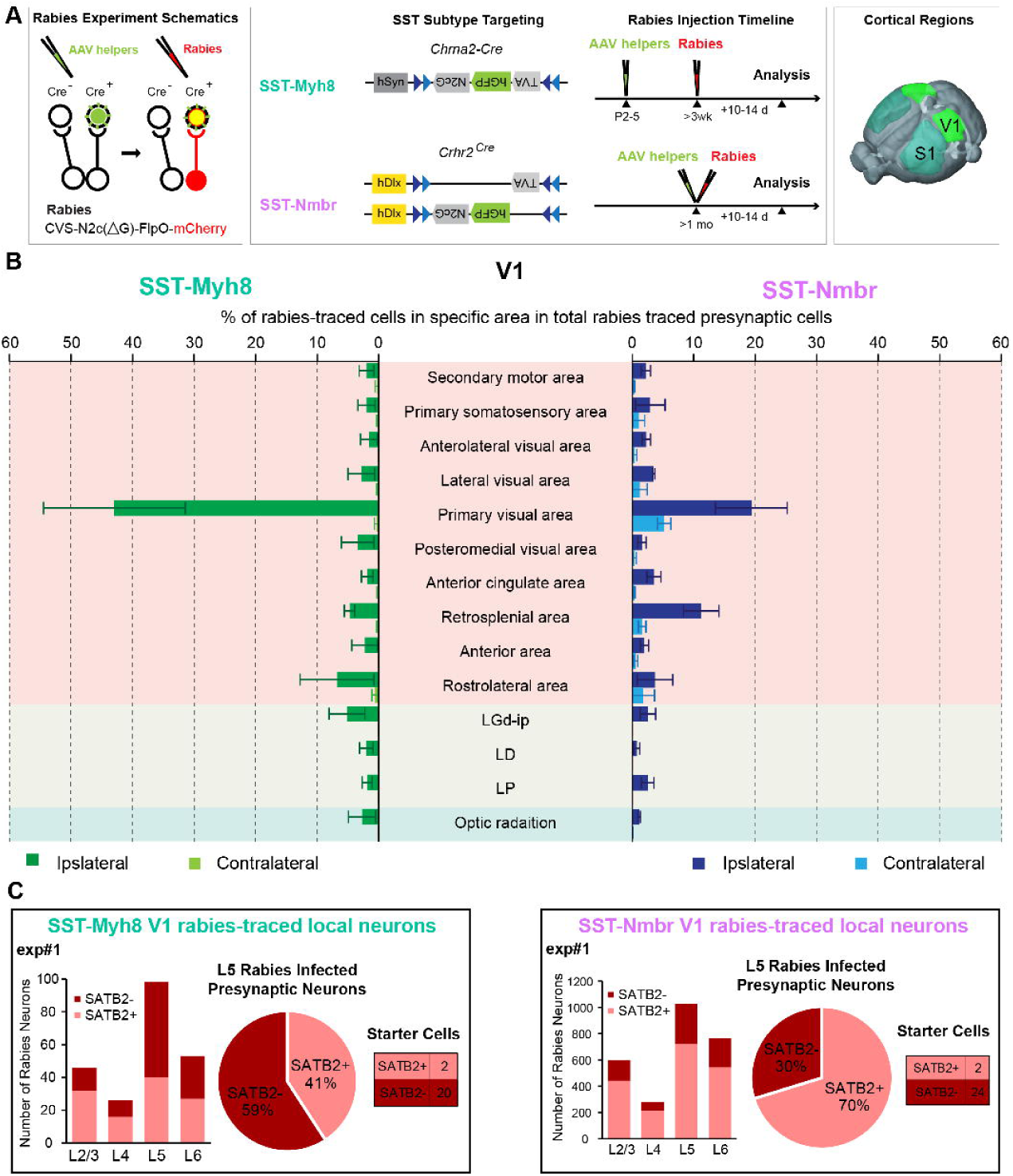
Monosynaptic rabies tracing from two different SST subtype revealed cell type specific afferent input. (A) Experimental design of rabies retrograde tracing from two SST subtypes. (left panel) Principle of rabies tracing TVA and N2cG conditional helpers (in green) are expressed using AAVs, followed by the specific infection and retrograde labeling by EnVA-pseudotyped CVS N2c rabies virus (Rabies, in red). (middle panel) The design of AAV-DIO-helper viruses and the timeline of AAV-helpers and N2cRV injections for tracing from SST-Myh8 (top) and SST-Nmbr interneurons (bottom) using *Chrna2-Cre* and *Crhr2^Cre^*mouse lines, respectively. Rabies tracing patterns were analyzed 10-14 days post-infection. (right panel) The tracing was performed on both SST subtypes from two cortical regions, S1 and V1. (B) Presynaptic inputs to SST-Myh8 and SST-Nmbr interneurons in V1 quantified as the percentage of rabies traced cells in each regional category out of the total number of cells labeled in the brain. Top 10 input regional categories for either SST subtype are included in the plot. Three rabies tracing experiments were performed for each SST subtype. (C) Quantification of rabies traced local presynaptic neurons in one representative experiment from SST-Myh8 (left) and SST-Nmbr interneurons (right), respectively. Immunostaining against SATB2 helps to reveal the identity of presynaptic neurons: SATB2+ neurons are IT neurons, SATB2-neurons are either PT neurons or interneurons. For each experiment, histogram of rabies traced neurons in each layer (left) is indicated. Pie chart (middle) shows the percentage of SATB2+ and SATB2-neurons labeled by rabies in L5, and table (right) shows the number of starter cells. As expected, most of the starter cells are SATB2-as expected, though occasionally there is small contamination of a low number of SATB2+ pyramidal neuron starter cells, which is inevitable since the targeting population only constitute ∼2% of cortical neurons. See also Figure S7-8.

We quantified the retrogradely labeled presynaptic neurons and normalized the number to the total amount of rabies traced presynaptic cells. Overall, both SST subtypes primarily received input from local excitatory neurons, other cortical regions, and the corresponding sensory thalamus relative to the site of injection (Figure 6B, Figure S7B-C) (Pouchelon et al., 2021). The top 10 brain regions for both SST subtypes combined, which contain almost exclusively cortical regions and thalamus, could account for >70% of all afferent inputs identified (Figure 6B, Figure S7C). As expected, the topmost afferent region for both SST subtypes is the injection area, suggesting that SST interneurons primarily receive inputs from local excitatory neurons. Intriguingly, one difference noted was that SST-Nmbr interneurons seemed to receive more inputs from contralateral cortex, while inputs to SST-Myh8 interneurons were almost exclusively from the ipsilateral side (n = 3 for SST-Myh8, n = 3 for SST-Nmbr in V1, Figure 6B; n = 3 for SST-Myh8, n = 2 for SST-Nmbr in S1; Figure S7C). Notably, different AAV-helper viruses were utilized for targeting SST-Myh8 and SST-Nmbr interneurons, and we observed many more rabies traced presynaptic neurons relative to the number of starter cells when tracing from SST-Nmbr interneurons as compared to SST-Myh8 interneurons (Figure S7B). As such, we wanted to confirm that the observed differences in contralateral versus ipsilateral inputs to these two SST subtypes was not an experimental artifact. Specifically, we suspected that the larger amount of retrogradely traced presynaptic neurons in SST-Nmbr experiments was likely due to a higher level of G protein expression in SST-Nmbr starter cells, which utilized a more efficient rAAV construct. We therefore repeated one rabies tracing experiment from SST-Myh8 interneurons in S1, using the same AAV-helper viruses used for targeting SST-Nmbr (Figure S7D). Reassuringly, this experiment revealed few contralateral inputs to SST-Myh8 interneurons, despite yielding a larger number of retrogradely traced cells. Given that SST-Nmbr preferentially targets L5-IT neurons, while SST-Myh8 primarily targets L5-PT neurons, the larger fraction of contralateral inputs to SST-Nmbr interneurons could reflect preferred afferent connectivity from IT neurons.

To examine whether these two SST subtypes received inputs from distinct populations of local pyramidal neurons, we performed immunostaining against SATB2, a marker for IT neurons in the mature cortex (McKenna et al., 2015), to determine the identity of the retrogradely traced local input neurons (Figure S8). The laminar distribution of the local inputs to both SST subtypes was very similar. The majority of the presynaptic neurons resided in the infragranular layers and most of them were found in L5 (Figure 6C, Figure S8). This result correlates with output mapping indicating that both SST subtypes preferentially target L5, despite SST-Nmbr interneurons residing primarily in L6 (Figure 3G). This also suggests that these two subtypes receive reciprocal innervations from L5 excitatory neurons. Furthermore, this distribution pattern is consistent between S1 and V1 (Figure S8), suggesting that the selective input and output connectivity we described might be stereotyped microcircuit properties intrinsic to different SST subtypes. Intriguingly, we found that the majority of L5 inputs to SST-Myh8 are SATB2-negative, indicating that they were either L5-PT neurons or interneurons (n = 4/5, S1 and V1 combined, Figure 6C and Figure S8). In contrast, the majority (70-80%) of L5 presynaptic neurons to SST-Nmbr interneurons are SATB2-positive, suggesting that they were L5-IT neurons (n = 4, S1 and V1 combined, Figure 6C and Figure S8). Again, this input connectivity seems to mirror the output connectivity, SST-Myh8 interneurons preferentially projected to L5-PT neurons, while SST-Nmbr interneurons preferentially connected to L5-IT neurons. This suggests that there are reciprocal selective connections between specific SST subtypes and excitatory neurons.

### Two Martinotti SST subtypes showed distinct subcellular innervation of L5 PT dendrites

Our results have demonstrated that SST subtypes showed selective input/output connectivity that is both laminar and cell type specific. Of particular interest are L5-PT neurons, which received strong input from both SST-Calb2 and SST-Myh8 interneurons (Figure 4). We therefore wondered whether input from these SST types was functionally redundant, or whether they provided qualitatively distinct forms of inhibition to a common target. Notably, these interneuron subtypes have distinct axonal morphologies: SST-Calb2 has a fanning-out shape, whereas SST-Myh8 has a T-shape (Figure 2). One possibility is therefore that they could impinge on different subcellular compartments of L5-PT dendrites. To test this, we quantified the distribution of synaptic puncta from SST-Calb2 and SST-Myh8 interneurons onto virally labeled L5-PT basal, oblique, apical branch, and tuft dendrites (Figure 7A-D). Immunostaining for both the presynaptic marker Gad65 and the postsynaptic marker Gephyrin allowed us to identify putative inhibitory synaptic boutons at the intersection of four fluorescent channels (see Methods). We found that SST-Calb2 puncta were distributed across the apical dendritic arbors, with the greatest density situated on the tuft and apical branches (Figure 7E,G). SST-Myh8 puncta, on the other hand, were concentrated solely on the tuft (Figure 7F). These results demonstrate that two SST subtypes impinging on the same excitatory population have distinct innervation patterns at a subcellular level, providing further support for the interesting possibility that they may gate different streams of information to a common target, as suggested in (Muñoz et al., 2017).

**Figure 7.**
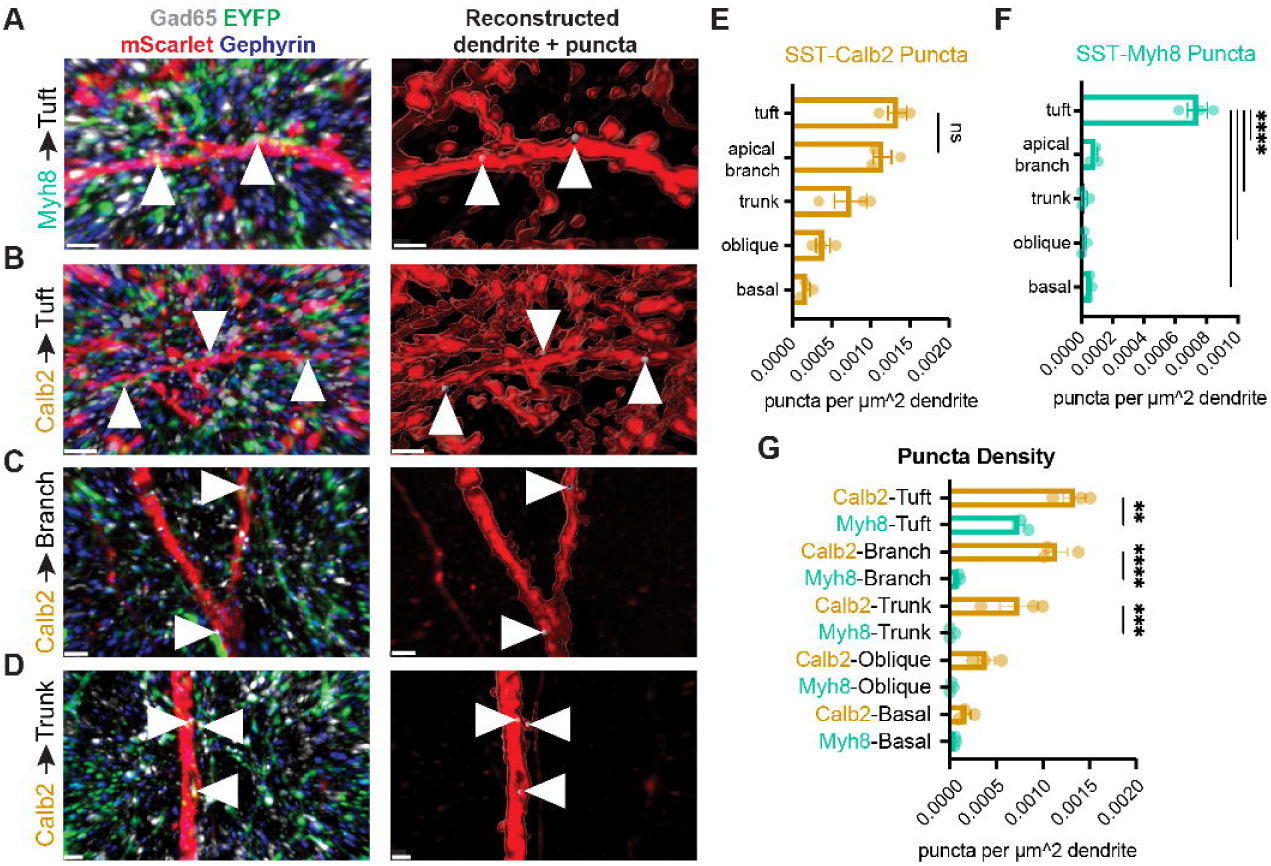
SST subtypes target distinct subcellular compartments of L5-PT dendrites. (A) Representative image of immunostaining (left) and reconstruction (right) of SST-Myh8 synaptic puncta on PT tuft dendrites. SST-Myh8 axons are labeled with Ai80-ChR2-EYFP, L5-PT dendrites labeled with rAAV2-retro-hSyn-mScarlet, presynaptic puncta labeled with Gad65, and postsynaptic puncta labeled with Gephyrin. Puncta (arrowheads) are the colocalization of all four channels. Reconstruction shows the dendrite surface and location of synaptic putative puncta. (B) As in (A) for SST-Calb2 puncta on L5-PT tuft dendrites. (C) As in (A) for SST-Calb2 puncta on a L5-PT dendritic apical branch. (D) As in (A) for SST-Calb2 puncta on L5-PT dendritic trunk. (E) Quantification of SST-Calb2 puncta on L5-PT dendrites. Number of puncta are normalized by the surface area of the reconstructed dendrite. Tuft and apical branch puncta densities were not significantly different (tuft vs. apical branch p = .2655) by one-way ANOVA with Tukey correction. All other comparisons revealed significantly more synapses on the tuft and apical branch than other compartments (tuft vs. trunk p = .0006, tuft vs. oblique p = .0009, tuft vs. basal p = .0002, apical branch vs. trunk p = .0135, apical branch vs. oblique p = .0195, apical branch vs. basal p = .0029). Comparisons between the remaining compartments were not significantly different. (F) Quantification of SST-Myh8 puncta on L5-PT dendrites. Number of puncta are normalized by the surface area of the reconstructed dendrite. Puncta density on tuft was significantly different for all comparisons (tuft vs. apical branch, tuft vs. trunk, tuft vs. oblique, and tuft vs. basal p < .0001) by one-way ANOVA with Tukey correction. (G) Comparison of SST-Calb2 and SST-Myh8 puncta on L5-PT dendrites. SST-Calb2 had significantly more puncta on the tuft (p = .001), apical branch (p < .0001) and trunk (p = .0001) than Chrna2 and a trend of more puncta on the oblique dendrites (p = .051) but not basal dendrites (p = .9101) by one-way ANOVA with Tukey correction.

## DISCUSSION

The rapid expansion of single-cell transcriptomic analysis of cell type taxonomies in recent years has resulted in an unprecedented understanding of the molecular heterogeneity of cortical neurons (Callaway et al., 2021; Ecker et al., 2017; Ngai, 2022). However, despite recent efforts (Berg et al., 2021; Bugeon et al., 2022; Cadwell et al., 2016; Condylis et al., 2022; Fuzik et al., 2016; Gouwens et al., 2020; Kim et al., 2020; Scala et al., 2021; Tasic et al., 2016, 2018), an understanding of how transcriptomic cell type correlates with other modalities including morphology, connectivity and *in vivo* functions are still largely lacking. In this study, we developed and characterized genetic strategies to target the breadth of transcriptomically identified SST subtypes. We then focused on three major SST subtypes and demonstrated that different transcriptomic subtypes form precise and partially reciprocal inhibitory microcircuits with excitatory and inhibitory neurons that are laminar, cell-type and subcellular specific. Previous studies also demonstrated that SST-Hpse and SST-Crh subtypes, form specific reciprocal connections with L4 spiny stellate cells (Naka et al., 2019; Nigro et al., 2018). A schematic diagram summarizing the characterized and hypothesized local microcircuitry formed by individual SST subtypes in S1 and V1 is shown in Figure 8. Therefore, taking SST interneurons as an exemplar, we provide a roadmap for understanding of interneuron subtypes, which emphasizes the previously underappreciated circuit specificity linking different subtypes of inhibitory and excitatory neurons.

**Figure 8.**
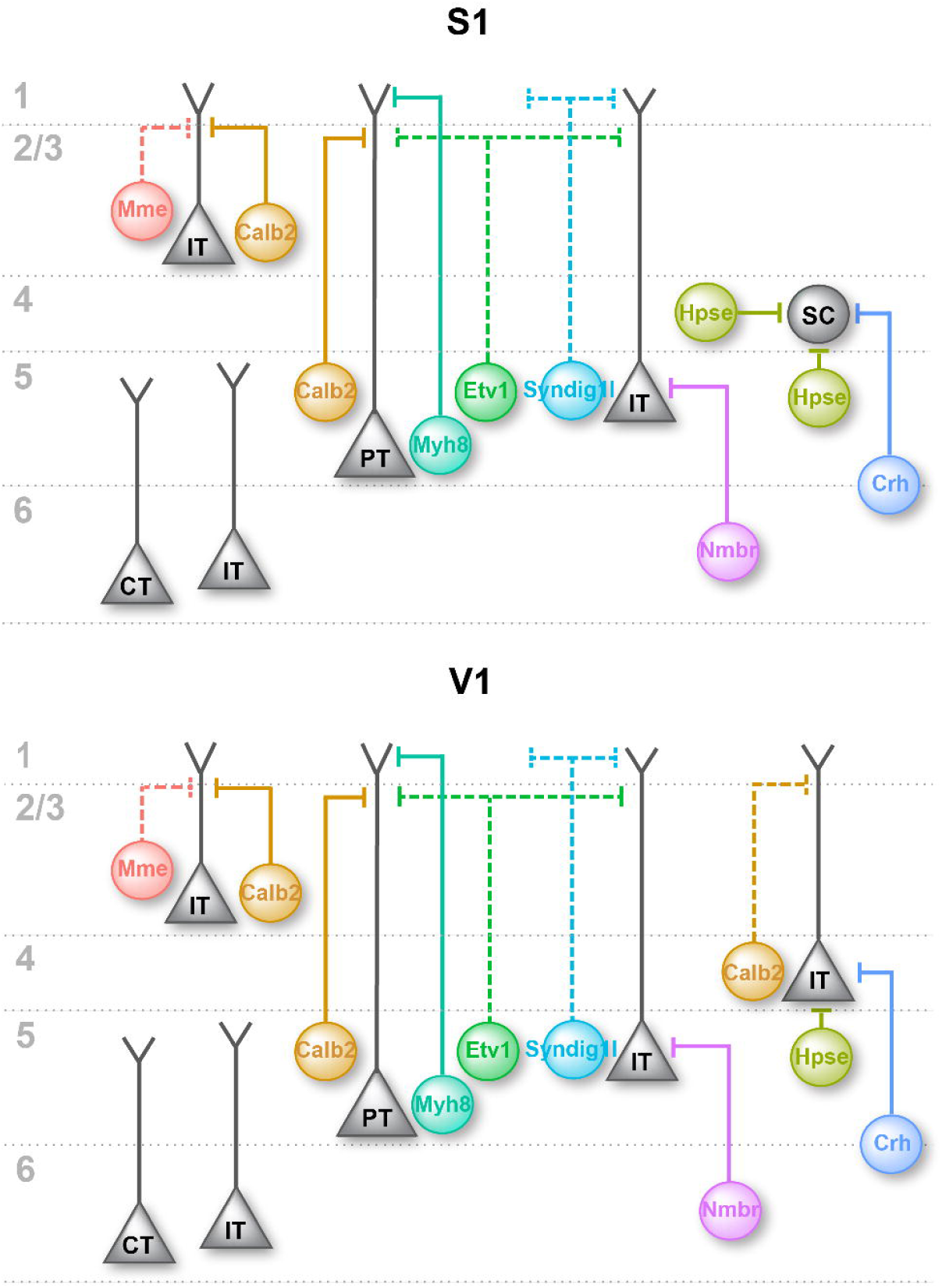
Schematic drawing of the output circuitry of different SST subtypes in S1 and V1. Summary of our current understanding about innervation pattern of different SST subtypes in S1 and V1, showing the preferred postsynaptic excitatory neuron cell type of each SST subtypes. Dashed lines are used for the output circuitry that are not fully characterized. IT, intratelencephalic neuron; PT, pyramidal-tract neuron; SC, L4 spiny stellate cell; CT, corticothalamic neuron.

### Inhibitory interneurons contribute to specific cortical microcircuits

Although the spatial distribution of cortical interneurons does not strictly obey the laminar boundaries set by excitatory neurons, our results demonstrate that, like the local excitatory network, cortical inhibitory circuits are organized in both a layer and cell-type specific fashion. Therefore, the complexity of inhibitory circuitry is at least as multifaceted as local excitatory neuron networks. A complete description of them requires knowledge of the laminar position of their afferent and efferent targets. While this study begins to characterize the local inhibitory microcircuits of SST subtypes with selected excitatory neuron types, the inclusion of other neuronal types such as L5 near-projecting pyramidal neurons, L6 corticothalamic neurons, and VIP interneurons will be necessary to gain a complete understanding of these inhibitory circuits. Nevertheless, our results provide a first-pass look at the granularity of their multilayered specificity.

Given this unanticipated specificity, upon reflection, it is not surprising that cortical inhibitory circuits were hypothesized to primarily exert ‘blanket’ inhibition when analyzed at the cardinal class level (Fino et al., 2013; Karnani et al., 2014). While SST interneurons as a class reside primarily in the infragranular layer, their overall inhibition to L2/3 excitatory neurons is equally as strong as to L5, resulting in a false impression of non-selective efferent targeting. By examining SST interneurons in terms of their different subtypes with respect to excitatory neurons, we observed that different layers of excitatory neurons receive inhibition roughly in proportion to the composition of SST subtypes found in their resident layer. Despite this, excitatory neurons clearly also receive a portion of SST inputs from populations residing in other layers (Yetman et al., 2019). For example, L5-IT neurons receive at least equally strong translaminar inhibition from the L6 SST-Nmbr subtype, as compared to SST-Myh8 and SST-Calb2 subtypes that both reside in L5. These results seem to suggest that the local inhibitory circuits have a hierarchical organization, whereby interneuron subtypes are distributed across layers to provide balanced laminar inhibition, while within each layer interneuron subtypes innervate specific postsynaptic inhibitory and excitatory neurons. The third layer of specificity is achieved subcellularly. SST-Myh8 and SST-Calb2 interneurons target L5-PT neurons with similar strength but innervate distinct dendritic domains. Thus, rather than inhibition being indiscriminate, specificity is achieved through the precise regulation of subtype number, subtype distribution, synaptic strength, and synaptic organization.

Appreciation of the fine structure of inhibitory circuitry provides the opportunity to rethink cortical function. As a more complete picture of the selectiveness of inhibitory circuitry emerges, the logic by which excitation is routed and gated will be forthcoming. Previously, The vast majority of work on SST interneurons has been done in the L2/3 supragranular regions where the SST-Calb2 population dominates (Adesnik et al., 2012; Fu et al., 2014; Gentet et al., 2012; Lee et al., 2013; Pi et al., 2013) However, the majority of SST interneuron diversity resides in the granular and particularly infragranular layers where SST interneuron diversity has been less explored, although see (Halabisky et al., 2006; Muñoz et al., 2017; Naka et al., 2019; Silberberg and Markram, 2007; Wang et al., 2004). These regions are also marked by a greater diversity of pyramidal cell classes. While supragranular layers are largely comprised of IT cells, the granular and infragranular cortices contain at least six distinct classes of pyramidal cells, including but not limited to IT, PT, near-projecting, corticostriatal, and corticothalamic neurons. Consistent with the specificity of inhibitory/excitatory circuits, we hypothesize that the diversity of interneurons scales with the diversity of pyramidal subtypes. We predict this allows class-specific pairing between different SST subtypes and excitatory populations.

Importantly, while excitatory neurons vary systematically across areal territories (Krienen et al., 2020), interneuron subtypes appear to be shared across the isocortex (Tasic et al., 2018). Nonetheless, based on the shared morphology and axonal projection patterns for the same SST subtype residing in two different cortical regions (S1 and V1), SST subtypes may function similarly across the isocortex. However, given the differences among related excitatory cell types across regions, it remains unclear how one SST subtype can adapt to homologous local excitatory neurons within distinct cortical areas. In this regard, the ability of two SST subtypes to target distinct dendritic domains of the same excitatory cell type (Figure 7), suggests more subtle biophysical control of inhibition. Given that differences in efferent specificity extend to the synaptic level, the modulation of long-range signals to specific pyramidal cells by different SST subtypes could be profound. An intriguing possibility is that these different SST subtypes target specific excitatory inputs, thereby gating distinct streams of information. As such, future exploration of dendritic integration, and the interactions between convergent inhibitory input are clearly warranted.

### Do SST subtypes receive reciprocal cell-type specific excitatory inputs?

This study has been focused on the efferent specificity of SST interneurons. This raises the question as to whether SST interneurons receive corresponding reciprocal afferent inputs from the local excitatory neurons that they target. Historically SST interneurons have been proposed to function to provide feedback inhibition to PT neurons (Kapfer et al., 2007; Silberberg and Markram, 2007). Our retrograde rabies tracing experiments seems to support this conclusion: SST-Myh8 interneurons both target and receive connections from L5-PT neurons, while SST-Nmbr interneurons preferentially reciprocally target L5-IT neurons. Consistent with our findings, SST-Myh8 interneurons in A1 appear to reciprocally form connections with thick-tufted and therefore PT L5 pyramidal neurons (Hilscher et al., 2017). Additional evidence supporting reciprocal connections comes from previous studies that demonstrated that L4-targeting non-Martinotti cells in L5 form selective and reciprocal connections with L4 spiny stellate cells in S1. In addition, L5 fanning-out Martinotti cells were found to primarily receive excitatory inputs from L2/3 in S1 and target L5 pyramidal neurons in S1 (Naka et al., 2019; Nigro et al., 2018). Our data suggest that SST-Calb2 interneurons, as a whole, strongly inhibit L5-PT pyramidal neurons and L2/3. Such translaminar inhibitory circuits have been described by previous studies, that are mediated through SST interneurons (Kapfer et al., 2007; Otsuka and Kawaguchi, 2009; Pluta et al., 2019).

Extrapolating from these findings, we hypothesize that molecularly diverse interneuron subtypes are embedded in highly specific circuit motifs that can be understood as functional units. This raises the possibility that sophisticated cortical neural networks exist comprised of combinations of computational modules, which can be understood as assemblies of distinct functional units, akin to those found in integrated circuits.

## AUTHOR CONTRIBUTIONS

S.J.W., E.S. and G.F. conceived the project and wrote the manuscript. S.J.W. and E.S. performed experiments and analyses. G.S. assisted with snRNA-seq and Slide-seq V2 experiment analysis. S.Y., L.A., D.H.C., A.S. assisted with smFISH experiments. M.S. assisted with smFISH analysis. S.Y. and Y.Q. assisted with single-neuron Neurolucida reconstruction. L.A. and Y.Q. assisted rabies tracing experiment analysis. D.R. performed one stereotaxic injection analysis. Q.X. produced some of the viruses used in the study. A.P. and B.R. provided biocytin filling images of SST-Myh8 interneuron. S.H. contributed to design of AAV helper virus. I.B., V.K., G.M., E.M. and F.C. assisted with Slide-seq V2 experiments. D.A.S. provided *Hpse^Cre^*mouse line.

## DATA AVAILABILITY

All data generated or analyzed during this study are included in the manuscript and supporting files. Additional sparse labeling images can be found at https://doi.org/10.7910/DVN/NQDIPG. Any additional data will be shared by the lead contact upon request.

## STAR*Methods

### RESOURCE AVAILABILITY

#### Lead contact

Further information and requests for reagent should be directed to the lead contact Gord Fishell

#### Materials availability

Plasmids and viruses created in this study are available upon request from the lead contact.

#### Data and code availability

Sparse labeling images of SST interneurons were deposited in Harvard Dataverse and can be accessed through the following link: https://doi.org/10.7910/DVN/NQDIPG

Scripts for clustering of snRNA-seq data and analysis of Slide-seq V2 experiments are available at https://github.com/gs512/slideseq-engine.

### EXPERIMENTAL MODEL AND SUBJECT DETAILS

#### Mice

All experiments were approved by and in accordance with Harvard Medical School IACUC protocol number IS00001269. Animals were group housed and maintained under standard, temperature-controlled laboratory conditions. Mice were kept on a 12:12 light/dark cycle and received water and food *ad libitum*. Both female and male animals were used in all experiments. Transgenic mouse lines used in this study is included in Key Resources Table.

### METHOD DETAILS

#### Tamoxifen Induction

Tamoxifen (Sigma-Aldrich, T5648) was dissolved in corn oil (Sigma-Aldrich) at 10 or 20 mg/ml) with agitation or sonication and stored at −80°C or at RT and used within one week of preparation. Tamoxifen solution was administrated to mice through oral gavage. A wide range of tamoxifen is administrated to achieve different level of recombination. To achieve sparse labeling of SST interneurons for examining single-neuron morphology, a single dose of 0.5 - 2 mg of tamoxifen was administrated to *Pdyn^CreER^; Ai14*, *Pdyn^CreER^; Ai32* or *Etv1^CreER^; Sst^FlpO^; RC::FPSit* mice . To induce higher level of recombination for assessing the specificity and coverage of different genetic targeting strategies, a varying dosage ranging from a single dose of 1 mg up to 5 doses of 2 mg of tamoxifen per mouse was administered to *Pdyn^CreER^; Ai14*, *Pdyn^CreER^; Npy^FlpO^; Ai9* and *Etv1^CreER^; Sst^FlpO^; Ai65* mice. All Tamoxifen administrations were performed on mice aged from 2-week to 3-month.

#### Perfusion and Immunohistochemistry

For all histological experiments, mice were deeply anesthetized with sodium pentobarbital (Euthasol) by intraperitoneal injection and transcardially perfused with 1X PBS followed by 4% paraformaldehyde (PFA) in 1X PBS. Brains were dissected out and post-fixed overnight at 4°C.

To examine the expression pattern of transgenic mouse lines, immunofluorescence is routinely used to amplify the fluorescent signal of reporter protein labeling. Fixed brain samples then cryopreserved in 30% sucrose in 1X PBS. 40 µm brain sections were obtained through a Leica sliding microtome. For immunofluorescence, free-floating brain sections were incubated in primary antibodies diluted in antibody incubation solution (5% normal donkey serum, 0.25% Triton X-100 in 1X PBS) in coldroom overnight or up to three days. Secondary antibodies were diluted in antibody incubation solution at RT for 1-3 hrs, or in coldroom overnight.

For sparse labeling and single neuron morphology reconstruction, fixed brain samples were sectioned through a vibratome (Leica VT1200S) into 100-150 µm slices. Brain sections were incubated in primary antibodies diluted in antibody incubation solution in coldroom for 2-3 days. Secondary antibodies were diluted in antibody incubation solution at RT for 1-3 hrs, or in coldroom overnight or up to 2 days.

For synaptic puncta staining, tissue was sectioned at 50 µm on a Leica VT 1200S vibratome. Free-floating brain sections were stored in antifreeze solution until processing. Free-floating brain sections were blocked for one hour (0.1% Triton X-100, 3% Normal Donkey Serum and 3% Normal Goat Serum in 1X PBS) for 1 hour, followed by incubation in the same solution overnight at 4°C. The following day sections were rinsed in 0.1% Triton X-100 in 1X PBS for a minimum of 3 x 5 minutes, followed by secondary incubation in the same blocking solution for 2 hours at room temperature. Sections were then rinsed again for a minimum of 3 x 5 minutes in 1X PBS and mounted.

A list of primary antibodies used in this study can be found in Key Resources Table.

#### Slide-seq V2

Slide-seq V2 experiments were performed on 10 μm thick coronal sections from four different wild-type mice aged between P28-37. Experimental procedures were detailed previously (Stickels et al., 2021). Samples were sequenced on an Illumina NovaSeq SP flow cell 100 cycle kit with 8 samples per run (four samples per lane). The Slide-seq tools (https://github.com/MacoskoLab/slideseq-tools) software was used to collect, demultiplex and sort reads across barcodes.

In addition, one published dataset from somatosensory cortex was included in the analysis, which can be accessed through https://singlecell.broadinstitute.org/single_cell/study/SCP815/sensitive-spatial-genome-wide-expression-profiling-at-cellular-resolution#study-summary.

#### Single Molecule Fluorescent In Situ Hybridization Histochemistry

For single molecule fluorescent in situ hybridization (smFISH) combined with immunohistochemistry, mice were perfused and brains were fixed overnight in 4% PFA in 1X PBS followed by cryoprotection in 30% sucrose in 1X PBS. Then, 16-20 μm (for RNAscope®) or 40-80 μm (for HCR-FISH) thick brain sections were obtained using a Leica cryostat or sliding microtome. Brain slices sectioned using a cryostat are directly mounted on glass slides (Fisherbrand Superfrost Plus) and preserved in -80 °C. Brain sections obtained using sliding microtome were preserved in Section Storage Buffer containing 28% (w/v) sucrose, 30% (v/v) ethylene glycol in 0.1M sodium phosphate buffer, pH 7.4, before smFISH experiments.

For RNAscope® experiments, samples were processed according to the ACDBio Multiplex Flourescent v2 Kit protocol (ACDBio #323100) for fixed frozen tissue. Briefly, tissue was pre-treated with a series of dehydration, H_2_O_2_, antigen retrieval and protease III steps before incubation with the probe for 2 hours at 40 °C. Note here protease III incubation was performed at room temperature to better preserve the protein for immunostaining. A list of probes purchased from ACDBio is included in Key Resources Table. Three amplification steps were carried out prior to developing the signal with Opal™ or TSA® Dyes (Akoya Biosciences). Immuostaining following RNAscope® experiment was performed according to Technical Note 323100-TNS from ACDBio. Samples were counterstained with DAPI and mounted using Prolong Gold antifade mounting medium (Molecular Probes #P369300).

HCR RNA-FISH experiments were performed with a modified protocol to manufacturer’s recommendation (Molecular Instruments). Briefly, three to four 40 or 80 µm brain slices were placed in a single well of a 24-well plate. The brain slices then went through a series of pre-treatment including post-fixation, an optical ethanol dehydration step, and a mild proteinase K treatment (2 µg/ml, 15 min, RT), before incubating with 3.3-4.5 nM of HCR RNA-FISH probes at 37 °C overnight. After repeated wash with probe wash buffer and 5X SSCT, the signal is developed and amplified with hairpin pairs at 60 nM at RT for 4-16 hrs. After amplification step, the brain slices were washed with 5X SSCT for 1.5 hr with periodic buffer change. Immunostaining following the HCR RNA-FISH was performed by blocking the brain slices with 2% BSA/PBST for ∼15 min, followed by overnight incubation with primary antibody diluted in 1% BSA/PBST at 4 °C overnight. After wash with 1X PBST, the brain slices are incubated with secondary antibodies diluted in 1% BSA/PBST at RT for 1-2 hrs. Brain slices were counterstained with DAPI (5 μM, Sigma #D9542) and mounted using Fluoromount-G (Invitrogen) or Prolong Gold antifade mounting medium (Molecular Probes #P369300). HCR RNA-FISH probes and amplifiers used in this study can be found in Key Resources Table.

#### Cell Culture, transfection and AAV production

HEK293FT cells (Thermo Fisher Scientific, #R70007) were cultured in Dulbecco’s Modified Eagle’s medium with high glucose and pyruvate, GlutaMAX Supplement, 10% fetal bovine serum, penicillin (100 units/ml) and streptomycin (100 μg/ml). The cultures were incubated at 37°C in a humidified atmosphere containing 5% CO2. For AAV production, HEK293FT cells were seeded on 15-cm plates without antibiotics for 24 hours and co-transfected with the following plasmids using Polyethylenimine (100 μg/dish, Polysciences, #23966-1): pHGTI-helper (22 μg/dish), rAAV2-retro helper (Addgene plasmid #81070, 12 μg/dish), AAV9 helper (Addgene plasmid #112865, 12 μg/dish),. and the AAV expression vector (12 μg/dish). 72 hours after transfection, transfected cells were harvested and lysed (150 mM NaCl, 20 mM Tris pH8.0) by three freeze-thaw cycles and Benzonase treatment (375 U/dish; Sigma, #E1014) for 15 minutes at 37°C. The supernatants were cleared by centrifugation at 4000 RPM for 20 minutes at 4 °C, then transferred to Iodixanol gradients (OptiPrep Density Gradient Medium, Sigma, #D1556) for ultracentrifugation (VTi50 rotor, Beckman Coulter) at 50,000 RPM for 1.5 hours at 16 °C. The 40% iodixanol fraction containing the AAVs was collected, underwent ultrafiltration with PBS in Amicon Ultra (15 ml, 100K, Millipore, #UFC910024) for 4 times, aliquoted and stored at −80 °C. The number of genomic viral copies was determined by qPCR using the following primers against the WPRE sequence: Fw: AGC TCC TTT CCG GGA CTT TC and Rv: CAC CAC GGA ATT GTC AGT GC. A list of viral vectors used in this study can be found in Key Resources Table.

#### Viral labeling of IT/PT neurons and PV interneurons

Juvenile mice (P10-15) were head-fixed using soft tissue Zygoma ear cups (Kopf #921). rAAV2-retro-hSyn-mScarlet (Dr. David Ginty) was used for retrograde labeling. Viral aliquots were loaded into a Drummond Nanoinjector III. All coordinates are referenced from lambda. For PT labeling, 150 nl was injected into the ipsilateral superior colliculus at AP 0.15, ML 0.38, DV - 1.45. For L5-IT labeling, 100 nl was injected into the ipsilateral retrosplenial cortex at AP 1.1, ML 0.39, DV -.25 and 50 nl at AP 0.5, ML 0.31, DV -.25. For L6-IT labeling, 150 nl was injected into contralateral V1 at AP 0.2, ML 2.0, DV -.45. For PV labeling, 200 nl of rAAV PHP.eB-S5E2-GFP-fGFP (Addgene #135631, Titer: 9.4x10^11^ vg/mL) was injected into ipsilateral V1 at AP .2, ML 2.0, DV -.45. Coordinates were slightly adjusted based on the age of the mouse at the time of injection (+/- .2).

#### Slice preparation and brain slice recording

Animals aged P25-31 were anesthetized with isoflurane followed by decapitation. The brain was quickly removed and immersed in ice-cold oxygenated sucrose cutting solution containing (in mM) 87 NaCl, 75 Sucrose, 2.5 KCl, 1.25 NaH_2_PO_4_, 26 NaHCO_3_, 10 Glucose, 1 CaCl_2_, 2 MgCl_2_ (pH=7.4). 300 µm thick coronal slices were cut using a Leica VT 1200S vibratome through primary visual cortex. Slices recovered in a holding chamber with ACSF containing (in mM) 124 NaCl, 20 Glucose, 3 KCl, 1.2 NaH_2_PO_4_, 26 NaHCO_3_, 2 CaCl_2_, 1 MgCl_2_ (pH=7.4) at 34 °C for 30 minutes and at room temperate for at least 45 minutes prior to recording.

For patch-clamp recording, brain slices were transferred to an upright microscope (Scientifica) with oblique illumination Olympus optics. Cells were visualized using a 60x water immersion objective. Slices were perfused with ACSF in a recording chamber at 2 ml/min at room temperature. All slice preparation and recording solutions were oxygenated with carbogen gas (95% O_2_, 5% CO_2_, pH 7.4). Patch electrodes (3–7 MΩ) were pulled from borosilicate class (1.5 mm OD, Harvard Apparatus). For current-clamp recordings patch pipettes were filled with an internal solution containing (in mM): 130 K-Gluconate, 10 KCl, 10 HEPES, 0.2 EGTA, 4 MgATP, 0.3 NaGTP, 5 Phosphocreatine and 0.4% biocytin, equilibrated with KOH CO_2_ to pH=7.3. For voltage-clamp recordings patch pipettes were filled with an internal solution containing (in mM): 125 Cs-gluconate, 2 CsCl, 10 HEPES, 1 EGTA, 4 MgATP, 0.3 Na-GTP, 8 Phosphocreatine-Tris, 1 QX-314-Cl, equilibrated with CsOH at pH=7.3. Recordings were performed using a Multiclamp 700B amplifier (Molecular Devices) and digitized using a Digidata 1550A and the Clampex 10 program suite (Molecular Devices). Voltage-clamp signals were filtered at 3 kHz and recorded with a sampling rate of 20 kHz. Recordings were performed at a holding potential of -70 mV. Cells were only accepted for analysis if the initial series resistance was less than 40 MΩ and did not change by more than 20% during the recording period. The series resistance was compensated at least ∼50% in voltage-clamp mode and no correction was made for the liquid junction potential. Experiments were performed at room temperature to ameliorate space clamp errors (Williams and Mitchell, 2008).

#### Optogenetic mapping

For output mapping, experiments were performed using mice express specific driver lines crossed with *Ai80* for intersectional CatCh expression and injected with AAVs to label IT, PT neurons, and PV interneurons. Whole-cell patch-clamp recordings were obtained from virally labeled neurons or unlabeled putative pyramidal neurons across layers. Virally labeled excitatory neurons were included as L5a and L5b neurons in the analysis of outputs across layers, but PV interneurons were excluded. For experiments with PV interneurons, at least 3 PV interneurons were included from each layer.

For optogenetic stimulation, 470 nm light was transmitted from a collimated LED (Mightex) attached to the epifluorescence port of the upright microscope. 1 ms pulses of light were directed to the slice in the recording chamber via a mirror coupled to the 60x objective (N.A. = 1.0). Flashes were delivered every 15 s over a total of 15 trials. The LED output was driven by a transistor-transistor logic output from the Clampex software.

#### Biocytin filling and staining

After recording with pipette solution containing 0.3-0.5% biocytin, the slices were fixed in 4% PFA overnight, then stored in 30% sucrose in 1X PBS till further processing. After washing out the PFA, the slices were incubated with *Scale*CUBIC-1 solution for 2 days. After thorough washing with 1X PBS, the slices were incubated with Alexa-conjugated streptavidin in blocking solution (10% normal donkey or goat serum, 0.5% Triton X-100, 0.2% cold water fish gelatin in 1X PBS) overnight at room temperature. After thorough wash with 1X PBS, slices were transferred to *Scale*CUBIC-2 solution and incubated for approximately 30 minutes before mounted on a glass slide in *Scale*CUBIC-2 solution for confocal microscopy imaging. Recipes for *Scale*CUBIC-1 and *Scale*CUBIC-2 can be found in (Susaki et al., 2014).

#### Retrograde monosynaptic rabies tracing

For tracing afferent inputs to SST-Myh8 subtype, *Chrna2-Cre* mouse pups at P2-5 were anesthetized by hypothermia and stereotaxically micro-injected with AAV9-DIO-helper virus encoding N2c-G-P2A-TVA-P2A-eGFP (Addgene #170853; Titer of 9.5x10^12^ vg/mL) using Nanoject III at a rate of 1 nL/s. AAV9-DIO-helper virus was diluted 1:1 or 1:2 with 1X PBS and injected for total of 10 nL (from Lambda: AP +1.5-2, ML +1.8-3, DV-0.2-0.3 for S1; AP +0-0.4, ML -1.8-2.2, DV -0.05-0.3 for V1). EnvA-pseudotyped CVS-N2c(ΔG)-FlpO-mCherry (N2c-RV, Titer: 3.7E+09U/ml) was generously shared by K. Ritola at Janelia Farms Research Center as described in (Pouchelon et al., 2021). N2c-RV was injected separately at P22-P42 (from Bregma: AP -1, ML +3, DV -0.85) for S1 or at P56-79 (from Bregma: AP -3, ML -2.5, DV -0.5) for V1. N2c-RV was diluted 1:10 with 1X PBS and stereotaxically injected for a total volume of 60-100 nL. Mice were sacrificed 10-15 days later for examination.

For tracing afferent inputs to SST-Nmbr subtype, *Crhr2^Cre^*mice at P34-79 were stereotaxically injected with AAV-Dlx-DIO-helpers and N2c-RV at the same time according to stereotaxic coordinates (from Bregma: AP −1, ML +3, DV-0.86 for S1; AP -3, ML -2.5, DV -0.50 for V1). AAV-Dlx-DIO-TVA, AAV-Dlx-DIO-GFP-N2cG and N2c-RV were combined in 1:1:1-3 ratio for injection of total 50-80 nL. For most of the rabies experiments, AAV9-Dlx-DIO-TVA (Titer: 6.89x10^13^ vg/mL) and AAV9-Dlx-DIO-GFP-N2cG (Titer: 5.46x10^13^ vg/mL) were used in combination, except for one experiment AAV1/2-Dlx-DIO-TVA (Titer: 3.5x10^12^ vg/mL) and AAV1/2-Dlx-DIO-GFP-N2cG (Titer: 2.9x10^12^ vg/mL) were used. Mice were sacrificed 13-14 days later for examination.

To confirm that the different afferent input pattern to these two SST subtypes were not caused by the use of different AAV-DIO-helper viruses, one test experiment was performed using AAV-Dlx-DIO-helpers from SST-Myh8 interneurons. Briefly, AAV9-Dlx-DIO-TVA and AAV9-Dlx-DIO-GFP-N2cG were stereotaxically injected in a P3 *Chrna2-Cre* mouse in 1:1 (v/v) ratio for a total of 10 nL (from Lambda: AP +1.8, ML +2.3, DV -0.25). N2c-RV was injected at P31. Mouse was sacrificed 15 days later for examination.

For all rabies tracing experiments, fixed brain samples were sectioned to 40 µm slices. One third of the total copy were taken for immunofluorescence experiments to examine the rabies tracing patterns.

#### Image acquisition

Images of transgenic mouse line labeling and rabies tracing were collected using a whole slide scanning microscope with a 10X objective (Olympus VS120 slide scanners) or using a motorized tiling scope (Zeiss Axio Imager A1) with a 5X or 10X objective.

Images of smFISH experiments were acquired with an upright confocal microscope (Zeiss LSM 800) with a 10X objective (Plan-Apochromat 10x/0.45 M27). For sparse labeling or biocytin filling experiments, imaged were acquired using the confocal microscope (Zeiss LSM 800) with a 20X objective (Plan-Apochromat 20x/0.8 M27). Images of synaptic puncta were acquired using an upright confocal microscope (Zeiss LSM800) with 40X oil immersion objective, 1.4 NA, 2.5 digital zoom, 1024 × 1024 pixels (∼0.22 μm resolution using 510 nm emission).

### QUANTIFICATION AND STATISTICAL ANALYSIS

Statistical details of experiments can be found in the figure legends. All statistical analyses were performed using GraphPad Prism or R software.

#### snRNA-seq pre-processing, clustering and label transfer

snRNA-seq datasets of interneurons in ALM and V1 using P28 Dlx5/6-Cre; Sun1-eGFP mice (here on referred to as Fishell_P28) were previously published (Allaway et al., 2021). Fishell_P28 and Allen mouse whole cortex and hippocampus Smart-seq dataset (https://portal.brain-map.org/atlases-and-data/rnaseq/mouse-whole-cortex-and-hippocampus-smart-seq) were pre-processed and aligned using Seurat (Satija lab). Supertype labels from the Allen dataset were transferred to Fishell_P28 dataset using Seurat integration. Briefly, Fishell_P28 was pre processed as described in (https://github.com/gs512/slideseq-engine/), with the clustering resolution set to 2.1.

Allen dataset was filtered using region_label, retaining cells from ALM, SSp, VISp, SSs and VIS regions. Cells were then split on class_label: GABAergic, Glutamatergic and Non-Neuronal. Glutamatergic cells were then filtered to retain cells with region_label equal to SSp and SSs; subclass_label CR, DG, L2/3 IT PPP, L5/6 IT TPE-ENT, L6b/CT ENT were excluded. Remaining cells were then processed in Seurat using default parameters (https://github.com/gs512/slideseq-engine/blob/82ceaebdca62d541dd044667f030fff5fb08bcea/helper.R#L70).

GABAergic cells were then filtered on supertype_label, excluding interneurons found outside of cortex: Meis2, Ntng1 HPF, Sst Ctsc HPF and Vip Cbln4 HPF. GABAergic interneurons in Allen dataset was integrated with Fishell_P28 using Seurat (https://github.com/gs512/slideseq-engine/blob/82ceaebdca62d541dd044667f030fff5fb08bcea/helper.R#L89) using default parameters, with the exception of dims 1:100 set for IntegrateData, and resolution = 1.7 for FindClusters. For each SST positive Integrated cluster, the most represented Allen supertype_label was transferred to the Fishell_P28 dataset.

#### Mapping SST subtypes from Allen Smart-seq dataset onto Slide-seq V2 with RCTD

The Allen mouse whole cortex and hippocampus Smart-seq dataset was processed as described above and used as the scRNA-seq reference for mapping SST subtypes in Slide-seq V2 data. We used the RCTD method (Cable et al., 2022), to integrate Allen Smart-seq data with spatial Slide-seq v2 data. Before running RCTD (now renamed as SPACEXR package), pucks were restricted to only include relevant cortical zones. SPACEXR was run in doublet mode, spots classified as doublet uncertain were not included in downstream analysis. Each non-excitatory spot was then assigned to a layer [L2/3, L4, L5, L6] using a KNN graph where the majority of the n nearest excitatory cells layer determined the assigned layer. Codes can be found on Github.

#### Quantification of marker gene expression and genetic labeling

Quantification of marker gene expression of smFISH experiments was performed by visual inspection of each genetically labeled neurons in maximum orthogonal projection images of confocal image stacks, either for all SST interneurons labeled by *Sst^Cre^;Ai14* or for neurons labeled by specific genetic strategies. Neurons contains at least three puncta were considered positive for the gene. In a few experiments where background noise was higher, the threshold was adjusted to five puncta per neuron. Quantification of calretinin-immunopositive neurons within *Etv1^CreER^;Sst^FlpO^;Ai65* labeled neurons was also performed by visual inspection.

For quantification of the percentage of genetic labeled neurons out of the total SST interneuron population, we performed smFISH against *Sst* mRNA for labeling all SST interneurons. Because *Sst* mRNA are abundant in SST interneurons and usually label the entire cell body, for majority of the analysis, we instituted a semi-automated strategy using Fiji (Image J), which achieved similar effectiveness as compared to a small subset of images that were analyzed by manual inspection. Briefly, the maximum orthogonal projection image of the Z-stack confocal image was loaded in FiJi program and the channel containing smFISH of *Sst* mRNA was selected. The image then went through a routine of brightness/contrast adjustment, background subtraction, smooth, and Gaussian Blur filtering before automatic ROI detection. Automatically detected ROIs were further selected based on size using Analyze Particles function in Fiji. An outline of all the selected ROIs were then superimposed over the original images, the final count of cell numbers was then performed manually using Cell Counter function in Fiji. A similar process is applied to the other channel containing genetically labeling neurons. ROIs from these two channels are then superimposed for identifying overlapping/non-overlapping neurons. DAPI channel was used to identify and quantify by cortical layer.

#### Neurolucida tracing

Z stacks of confocal images were loaded into Neurolucida 360 (MBF Biosciences). Cell body, dendrite and axons were recognized and reconstructed in a semi-automated manner. Neuronal processes were tracing using the ‘user guided’ option with Directional Kernels.

#### Intrinsic property recording and analysis

Passive and active membrane properties were measured in current-clamp from a holding potential of -65 mV. Analysis was done in Clampfit 11 and Prism (GraphPad). The electrophysiological parameters were adapted (Nigro et al., 2018) and defined as follows:

<u>Resting membrane potential (Vrest, in mV)</u>: membrane potential measured with no current applied, measured immediately after breaking into the cell;

<u>Input resistance (IR; in MΩ)</u>: resistance measured from Ohm’s law from the peak of voltage responses to hyperpolarizing current injections (up to −40 or −50 pA);

<u>Sag ratio (dimensionless)</u>: the ratio of voltage at the peak and voltage at steady-state in response to hyperpolarizing current injections, with the peak at approximately -90 mV;

<u>AP threshold (AP_thre_, in mV)</u>: measured from action potentials (APs) evoked at rheobase with 1 second current injections, as the membrane potential where the rise of the AP was 10 mV/ms;

<u>AP amplitude (mV)</u>: The peak amplitude measured from AP_thre_;

<u>AP half-width (in ms)</u>: duration of the AP at half-amplitude from AP_thre_;

<u>AP maximum rate of rise (AP_rise_, in mV/ms)</u>: measured from APs evoked at rheobase as the maximal voltage slope during the upstroke of the AP;

<u>After hyperpolarization potential (AHP, in mV)</u>: measured as the difference between AP_thre_ and the peak of the fAHP.

<u>Maximal firing frequency (HFF, in Hz)</u>: maximal firing frequency evoked with 1-s-long depolarizing current steps;

<u>Adaptation (dimensionless)</u>: measured from trains of approximately 35 APs as [1 − (Ffirst/Flast)], where Ffirst and Flast are, respectively, the frequencies of the first and last ISI; <u>Rebound APs</u>: number of APs elicited in response the end of a 1-s-long hyperpolarizing voltage deflection where the steady-state voltage response was approximately −90 mV.

#### Optogenetic experiment analysis

Data analysis was performed off-line using the Clampfit module of pClamp (Molecular Devices) and Prism 9 (GraphPad). Individual waveforms from 15 trials per cell were averaged, and the averaged peak amplitude was recorded. Data was tested for normality and lognormality in Prism and determined that in all cases the data was lognormally distributed (>90% likelihood). Kolmogorov-Smirnoff tests and Kruskal-Wallis tests with Dunn’s correction were used to determine statistical significance for experiments with single and multiple comparisons, respectively. To visualize the distribution of the data, for each graph the highest 75^th^ percentile value of all plotted datasets was selected as a breakpoint for the y-axis, with 75% of the data below the breakpoint and 25% above the breakpoint.

#### Hierarchical bootstrapping and pan-SST response simulation

Analysis was done in Matlab. We addressed two levels of variability in our data: the variability of single-cell IPSCs across 15 sweeps of ChR2 stimulation, and the variability across cells within a given condition (layer or pyramidal cell type). First, we recomputed the average IPSC amplitude per cell by sampling with replacement across all 15 sweep amplitudes and taking a new average. Second, we recomputed the set of IPSC amplitudes per condition by sampling from the set of bootstrapped amplitudes with replacement. To simulate the linear combination of three SST subtypes we added together one randomly selected one amplitude per subtype (SST-Calb2, SST-Myh8, and SST-Nmbr). We repeated the bootstrapping procedure for the pan-SST IPSCs and subtracted the simulated linear combination IPSC from the median pan-SST IPSC to compare the two amplitudes (for example an difference of 0 means the amplitudes of the combined SST-Calb2, SST-Myh8, and SST-Nmbr equaled the pan-SST response, and a difference > 0 means that the combined SST-Calb2, SST-Myh8, and SST-Nmbr response was greater than the pan-SST response). This was repeated 100,000 times to determine the distribution of the difference between the simulated and measured pan-SST response.

#### Monosynaptic rabies tracing analysis

Upon uploading all the images into NeuroInfo software, all sections were manually reordered from rostral to caudal of the brain. The software’s section detection parameters were adjusted to correctly recognize the borders of each brain section. Sections were aligned, first using the software’s Most Accurate alignment option, and adjusted manually if necessary. After specifying the distance (120 µm) between each section, the Section Registration function of the software would compare each section to an existing 3D model of the mouse brain to estimate the rostral-caudally location of each section. Non-linear registration was run on each section to account for the slight distortions that might happen during sectioning/mounting, and/or imperfections in sectioning angle. In the Cell Detection function, parameters for cell size and distance from background were adjusted, and then Neural Network with preset pyramidal-p64-c1-v15.pbx was used to automatically detect rabies infected cells in the red channel. Detection results were reviewed manually to correct for any detection mistakes (false positives or negatives). Starter cells were manually marked and identified as GFP co-localized rabies infected cells. Final results were exported to an Excel file for further analysis.

#### Synaptic puncta analysis

For analysis of synaptic density on dendritic compartments, the dendritic compartment was noted prior to image acquisition though in some cases a single image contained multiple compartments, requiring post-hoc segmentation.

To quantify the density of synaptic puncta on dendrites, 5 µm-thick image stacks were analyzed with IMARIS 9.5.0 or 9.7.0 using MATLAB scripts adapted from a previous study (Favuzzi et al., 2019). First all channels underwent background subtraction and depth normalization. Then three-dimensional “surfaces” using the “Create Surface” tool were automatically reconstructed for mScarlet+ dendrites and GFP+ SST cells with a 2 µm^2^ area filter. The threshold was selected to include as much of the process as possible while minimizing background noise. Surfaces were manually edited to exclude artifacts, segment dendritic compartments, and remove the soma and dendrites of SST interneurons when necessary (clearly distinguishable from axons by their size and brightness). Gad65+ and Gephyrin+ puncta were automatically reconstructed as “spots” of 0.6 and 0.3 µm diameter, respectively. To detect spots the built-in spot detection algorithm in Imaris first applies a 3D Mexican Hat filter using the spot size and then locates the spot centroid at the local maxima of the filtered image. Gad65+ spots located within the axon surface were identified using the “split into surface” tool using the radius of the spot as the threshold distance, and the same was done for Gephyrin+ spots in the dendrite surface. Finally, the presynaptic and postsynaptic boutons identified in the previous step were colocalized using their radii as a threshold to identify the number of puncta per image. That number was then normalized by the surface area of the reconstructed dendrite in that image.

To control for noisy signal, we reflected the Gad65 channel on the y-axis and repeated our analysis to determine how many puncta are detected by chance without the biologically correlated signal (Sup. Fig. 8A). As SST-Calb2 puncta were distributed across the dendritic arbor we included all images, but for SST-Myh8 interneurons we only included images of tuft dendrites as the other images contained little to no puncta. We found that for both SST-Calb2 and SST-Myh8, our analysis detected significantly more puncta in the original images (data not shown), confirming that we can detect synaptic puncta above a noise threshold.

## ACKNOWLEDGEMENT

We thank Marian Fernandez-Otero and Nusrath Yusuf and for technical assistance. Dr. David Ginty and Dr. Mark Springel for sharing with us the viral construct of pAAV-hSyn-mScarlet-WPRE. N2c-RV were generously shared by K. Ritola at Janelia Farms Research Center. This work was supported by grants from the National Institutes of Health (NIH)

R01NS081297, R37MH071679, and UG3MH120096 to G.F., the Simons Foundation SFARI to G.F., the Leonard and Isabelle Goldenson fellowship (FY19) to S.J.W., the William Randolph Hearst Fund (FY20) to S.J.W., F32 fellowship from National Institute of Mental Health F32MH125464 to S.J.W., F31 fellowship from National Institute of Neurological Disorders and Stroke F31NS110120 to E.S., and the BRAIN Initiative grant U19MH114830 to D.A.S.. We thank the Neurobiology Department and the Neurobiology Imaging Facility for consultation and instrument availability that supported this work. Finally, we are grateful for the comments and suggests of Drs. Chris Harvey, Emilia Favuzzi, Elisabetta Furlanis, Leena Ibrahim, as well as Areil Hairston. This facility is supported in part by the Neural Imaging Center as part of an NINDS P30 Core Center grant #NS072030.

