## SupplementalInformation for "Cortical somatostatin interneuron subtypes form cell-type specific circuits"

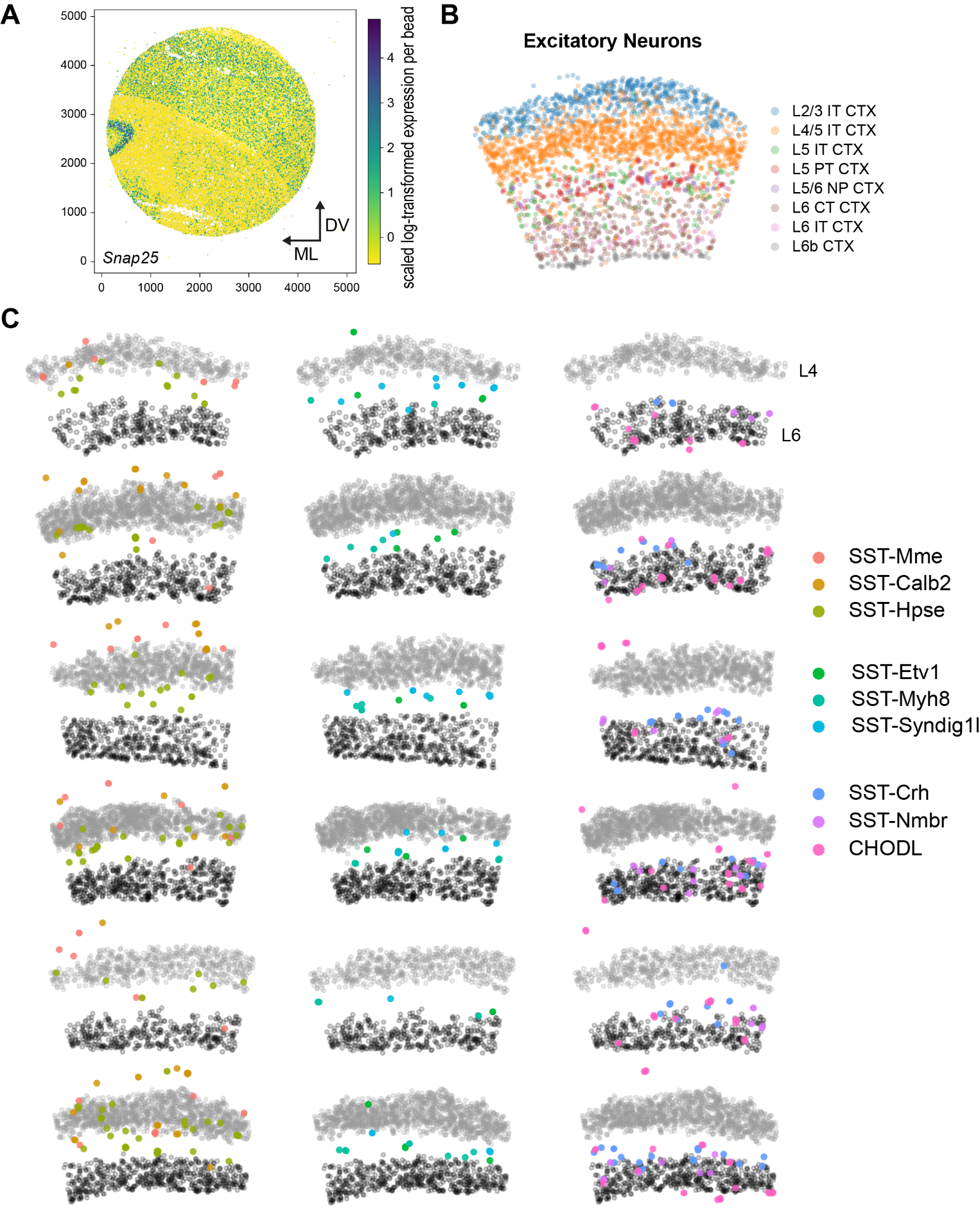

**Figure S1. Slide-seq V2 method: identification of cortical region, excitatory neuron assignment and SST subtype distribution in the rest six pucks. Related to STAR Method and Figure 1.**

(A) Expression of *Snap25* gene can assist in defining the cortex region in a coronal section of mouse brain processed by Slide-seq V2 (same experiment as shown in Figure 1C). Each dots represents a bead colored according to the scaled log-transformed expression level of *Snap25* gene. Subsequent analysis is restricted to cortical region where all cortical layers are represented (highlighted region).

(B) Same cortical region from (A), with dots colored by their predicted identity according to the assignment by robust cell type decomposition (RCTD). Only excitatory neurons are shown.

(C) The rest six Slide-seq V2 experiments in S1 region with RCTD predicted SST interneurons labeled. Grey circles showing the location of L4 and L6 excitatory neurons for reference.

| **Probe Gene Names** | ***Calb2*** | ***Hpse*** | ***Cbln4*** | ***Pdyn*** | ***Crh*** | ***Chodl*** |
| --- | --- | --- | --- | --- | --- | --- |
| **High Expression In** | SST-Calb2 CHODL | SST-Hpse | SST-Calb2 SST-Hpse | SST-Syndig1l SST-Hpse | SST-Crh | CHODL |
| **Low Expression In** | SST-Mme | SST-Syndig1l | SST-Mme | SST-Calb2  SST-Nmbr | SST-Nmbr |  |

**Table S1. List of smFISH probes and their expression levels in different SST subtypes. Related to Figure 1.**

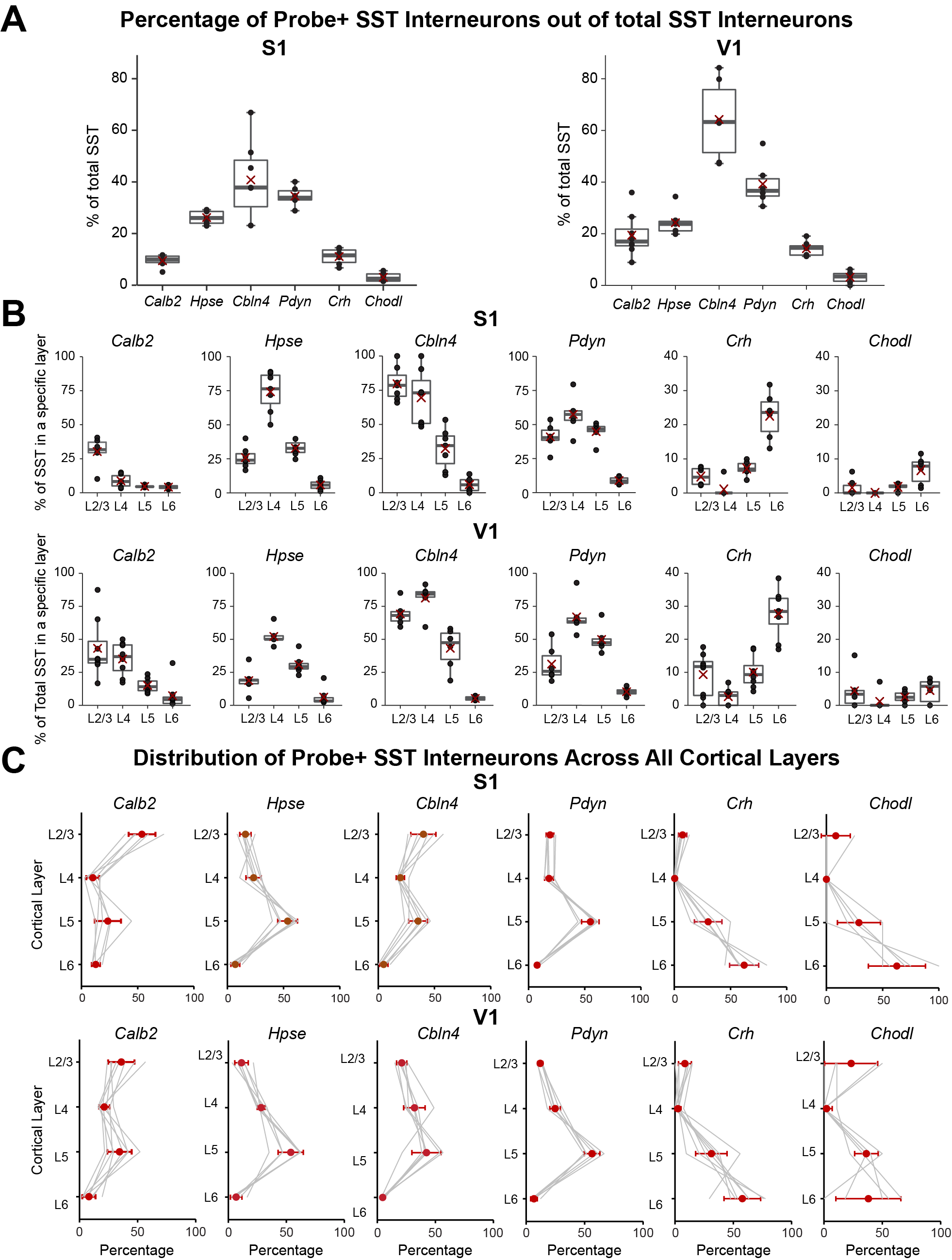

**Figure S2. Quantification of smFISH experiments against marker genes for various SST subtypes. Related to Figure 1.**

smFISH experiments against *Calb2*, *Hpse*, *Cbln4*, *Pdyn*, *Crh* and *Chodl* mRNA were performed on 1-2 month old *Sst^Cre^;Ai14* mice, where SST interneurons are labeled genetically and can be visualized by endogenous fluorescence. For all quantifications, at least three experiments performed on three different mice were used.

(A). Percentage of total SST interneurons in S1 (left) and V1 (right) that express specific marker genes as identified in smFISH experiments.

(B). Percentage of SST interneurons expressing specific marker genes in each layer in S1 (upper row) and V1 (lower row), as identified in smFISH experiments.

(C) The laminar distribution of SST interneurons expressing specific marker gene in S1 (upper row) and V1 (lower row).

| **SST Subtype** | **Cell#** | **Percentage** |
| --- | --- | --- |
| SST-Mme | 196 | 12.55% |
| SST-Calb2 | 307 | 19.65% |
| SST-Hpse | 230 | 14.72% |
| SST-Etv1 | 41 | 2.62% |
| SST-Myh8 | 221 | 14.15% |
| SST-Syndig1l | 60 | 3.84% |
| SST-Crh | 232 | 14.85% |
| SST-Nmbr | 208 | 13.32% |
| CHODL | 67 | 4.29% |

**Table S2. Proportions of different SST subtypes in the snRNA-seq dataset of P28 cortical interneurons in V1. Related to Figure 1.**

| **SST Subtypes** | **Genetic Strategy** | **Boolean Logic** |
| --- | --- | --- |
| **CHODL** | *Nos1^CreER^; Sst^FlpO^* | Cre AND Flp |
| **SST-Calb2**  **CHODL** | *Calb2^Cre^; Sst^FlpO^* | Cre AND Flp |
| **SST-Etv1**  **SST-Mme**  **SST-Calb2*** | *Etv1^CreER^; Sst^FlpO^* | Cre AND Flp |
| **SST-Myh8** | *Chrna2-Cre* | Cre only |
| **SST-Hpse**  **SST-Calb2** | *Pdyn^Cre^; Npy^FlpO^* | Cre AND Flp |
| **SST-Syndig1l**  **SST-Hpse*** | *Pdyn^CreER^* | Cre only**^†^** |
| **SST-Hpse**  **SST-Syndig1l** | *Hpse^Cre^* | Cre only**^†^** |
| **SST-Syndig1l** | *Pdyn^CreER^; Npy^FlpO^* | Cre-ON/Flp-OFF |
| **SST-Syndig1l** | *Pdyn^Cre^; Npy^FlpO^* | Cre-ON/Flp-OFF**^†^** |
| **SST-Crh** | *Crh^Cre^; Sst^FlpO^* | Cre AND Flp |
| **SST-Nmbr** | *Crhr2^Cre^; Sst^FlpO^* | Cre AND Flp |
| **CHODL**  **SST-Nmbr**  **SST-Mme** | *Tac1^Cre^; Sst^FlpO^* | Cre AND Flp |

**Table S3. Genetic strategies targeting different SST subtypes. Related to Figure 2.**

For genetic strategies that target multiple SST subtypes, the list of SST subtypes was arranged with primary target on the top and minor target at the bottom. * indicates that the SST subtype were labeled in variable degrees depending on the extent of Cre recombination. † indicates genetic strategies that work only at certain age range. *Hpse^Cre^* shows germline recombination. *Pdyn^Cre^* is expressed in a subset of excitatory neurons during development. *Tac1^Cre^* is expressed in CHODL interneurons early during development.

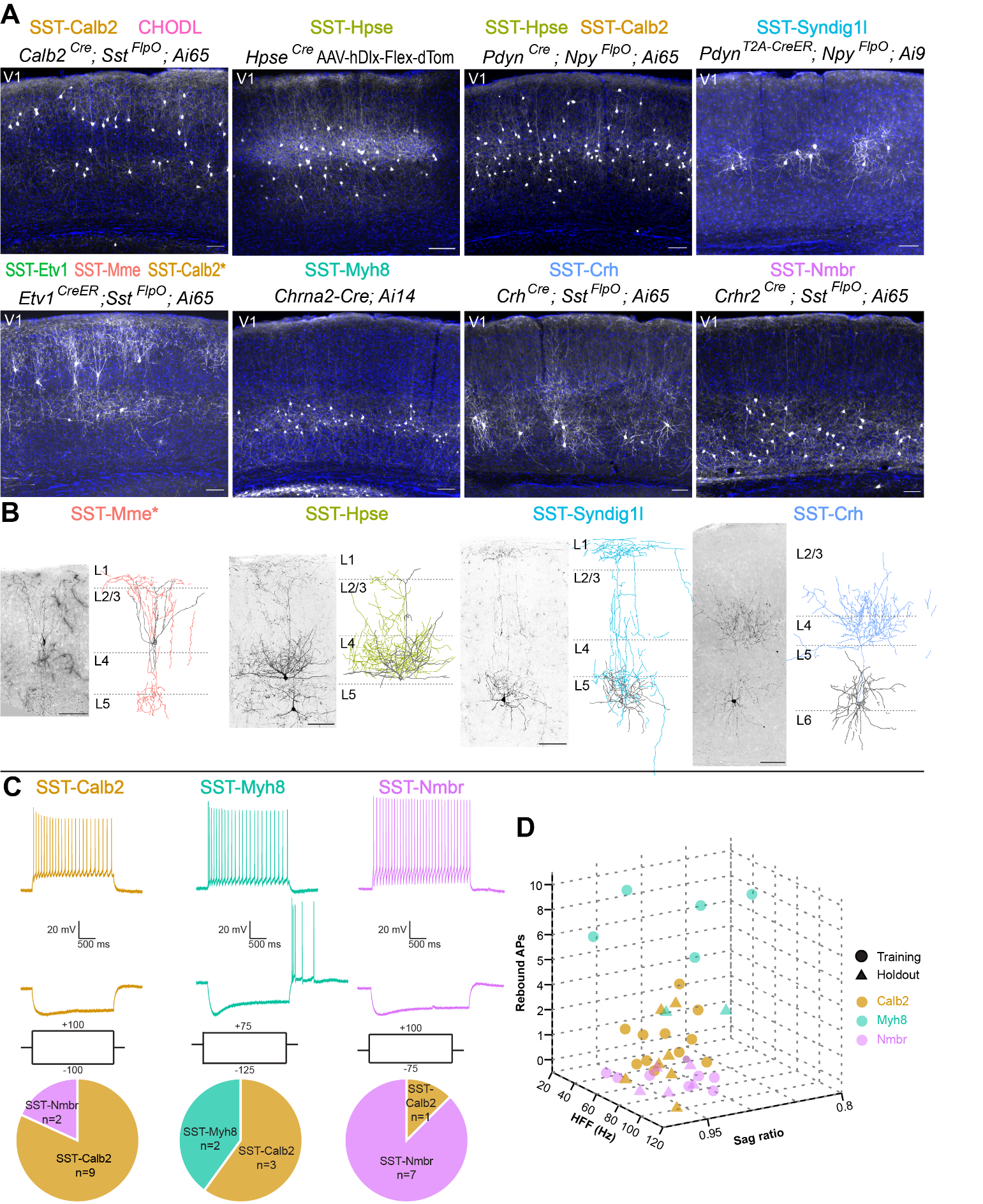

**Figure S3. Genetic labeling and single-neuron reconstruction of selected SST subtypes in V1; intrinsic electrophysiological properties of three SST subtypes in S1. Related to Figure 2.**

(A) Representative images of selective genetic strategies targeting SST subtypes in V1, with DAPI for laminar distribution. All images were taken from 1-3 month old mice. *Ai9* reporter line is used here as a Cre-ON/Flp-OFF strategy due to the FRT sites flanking the mutation that are still retained in this mouse line (Madisen et al., 2010). With this reporter line, SST-Hpse interneurons were occasionally observed in *Pdyn^T2A-CreER^; Npy^FlpO^; Ai9* strategy, likely due to incomplete FlpO recombination, though not noted in this representative image. For SST-Hpse labeling, rAAV9-hDlx-Flex-dTomato virus was stereotaxically injected in *Hpse^Cre^* mice in V1 at 1-month-old and examined 13 days post-injection. Scale bars, 100 µm.

(B) Sparse labeling of selective SST subtypes in V1. Images of genetic labeling is shown to the left of the Neurolucida reconstruction of single-neuron morphology. *Note that the example of SST-Mme sparse labeling was targeted by the intersectional strategy of *Etv1^CreER^; Sst^FlpO^;* *RC::FPSit*, which could also label SST-Etv1 and a varying degree of SST-Calb2. Because this example neuron resides in L2/3, whereas SST-Etv1 interneurons are expected to primarily reside in L5a, and that the chance of labeling SST-Calb2 is relatively low due to the low level of recombination, we infer that the identity of this labeled neuron is most likely SST-Mme. SST-Hpse and SST-Syndig1l interneurons are both labeled by *Pdyn^T2A-CreER^; Ai14* strategy and differentiated by their unique morphology. SST-Crh interneuron is labeled by *Crh^Cre^; Sst^FlpO^;RC::FPSit*. All reconstructions were derived using sparse labeling of specific SST subtypes from P25-73 mice. Scale bars, 100 µm.

(C) Representative traces of three SST subtypes in response to current injections. SST-Calb2, SST-Myh8, and SST-Nmbr all showed regular-spiking adapting firing patterns (top). Pie charts showing the number of cells classified as SST-Calb2, SST-Myh8, and SST-Nmbr interneurons by a trained k-nearest neighbor classifier (bottom, from left to right).

(D) 3D plot of the three most predictive features in the nearest neighbor analysis.

| **SST Subtype** | **Adult Marker Genes** | **Resident Layer** | **Axonal Project Pattern** | **Putative Cell Type** | **Tasic *et al.*, 2018 clusters** |
| --- | --- | --- | --- | --- | --- |
| **SST-Mme** | *Tac1, Necab1, Fam43a* | L2/3 | L1, L2/3 | Martinotti cell | Sst Tac1 Tacr3 Sst Tac1 Htr1d Sst Mme Fam114a1 |
| **SST-Calb2** | *Pcsk5* | L2/3, L5a (S1) L2/3, L4, L5a (V1) | L1, L2/3 | Fanning-out Martinotti cell | Sst Calb2 Pdlim5 Sst Calb2 Necab1 |
| **SST-Hpse** | *Ascl2* | L4, L5a | L4 | L4-targeting non-Marinotti cell | Sst Hpse Cbln4 |
| **SST-Etv1** | *Mstn* | L5 | L1, L2/3 | Martinotti cell | Sst Nr2f2 Necab1 Sst Myh8 Etv1 |
| **SST-Myh8** | *Chrna2, Plpp4, Cartpt, Myh13, Glra3* | L5b | L1, L5 | T-shaped Martinotti cell | Sst Chrna2 Glra3 Sst Myh8 Fibin Sst Myh8 Etv1 Sst Chrna2 Ptgdr |
| **SST-Syndig1l** | *C1qtnf7, Pdyn, Prdm1* | L5a | L1 | T-shaped Martinotti cell | Sst Hpse Sema3c Sst Chrna2 Ptgdr |
| **SST-Crh** | *Rxfp1, Prdm8, Ptprk, Lmo1* | L5b, L6 | L4, L5/6 | L4-targeting non-Marinotti cell | Sst Rxfp1 Prdm8 Sst Rxfp1 Eya1 Sst Tac2 Tacstd2 |
| **SST-Nmbr** | *Lpar1, Esm1, Crhr2* | L6 | L5/6 | L5/6-targeting non-Martinotti cell | Sst Crh 4930553C11Rik Sst Crhr2 Efemp1 Sst Esm1 Sst Tac2 Myh4 |
| **CHODL** | *Nos1, Sfrp1, Ntn1, Rasgef1b, Carhsp1* | L6 | L6, long-range | *Nos1^+^* non-Martinotti, long-range projecting neuron | Sst Chodl |

**Table S4. Summary of the current understanding about different SST subtypes. Related to Figure 2.**

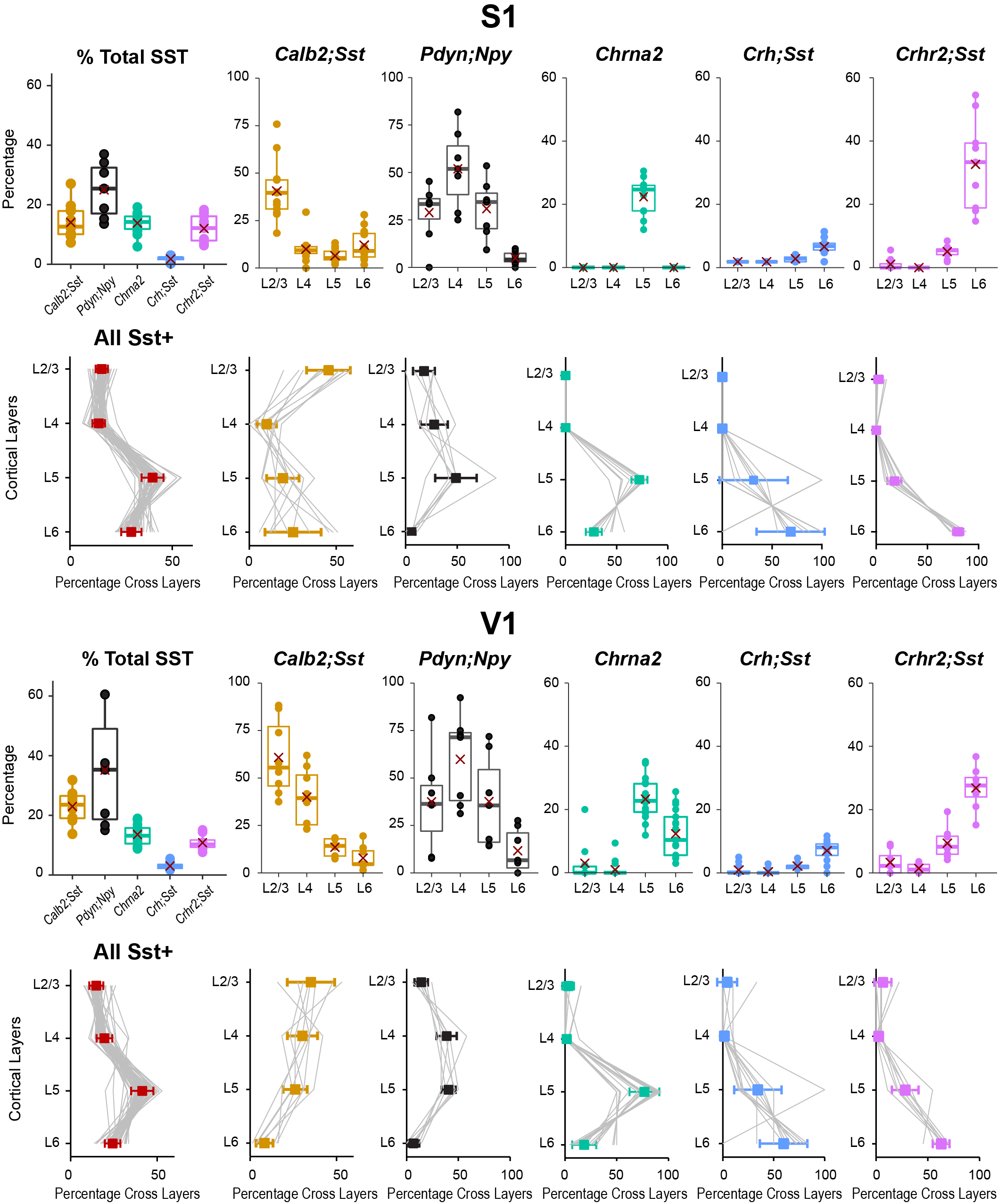

**Figure S4. Proportion and distribution of SST subtypes across cortical layers in S1 and V1. Related to Figure 2.**

All SST interneurons were labeled with smFISH against *Sst* mRNA. At least three experiments performed on three different mice were used for quantification.

(S1, top row) Proportion of genetically labeled neurons out of total SST interneurons (left most) or SST interneurons in each layer (rest) in S1.

(S1, second row) Laminar distribution of all SST interneurons (left most) or genetically labelled neurons by each strategy (rest) in S1.

(V1) Parallel quantifications performed in V1.

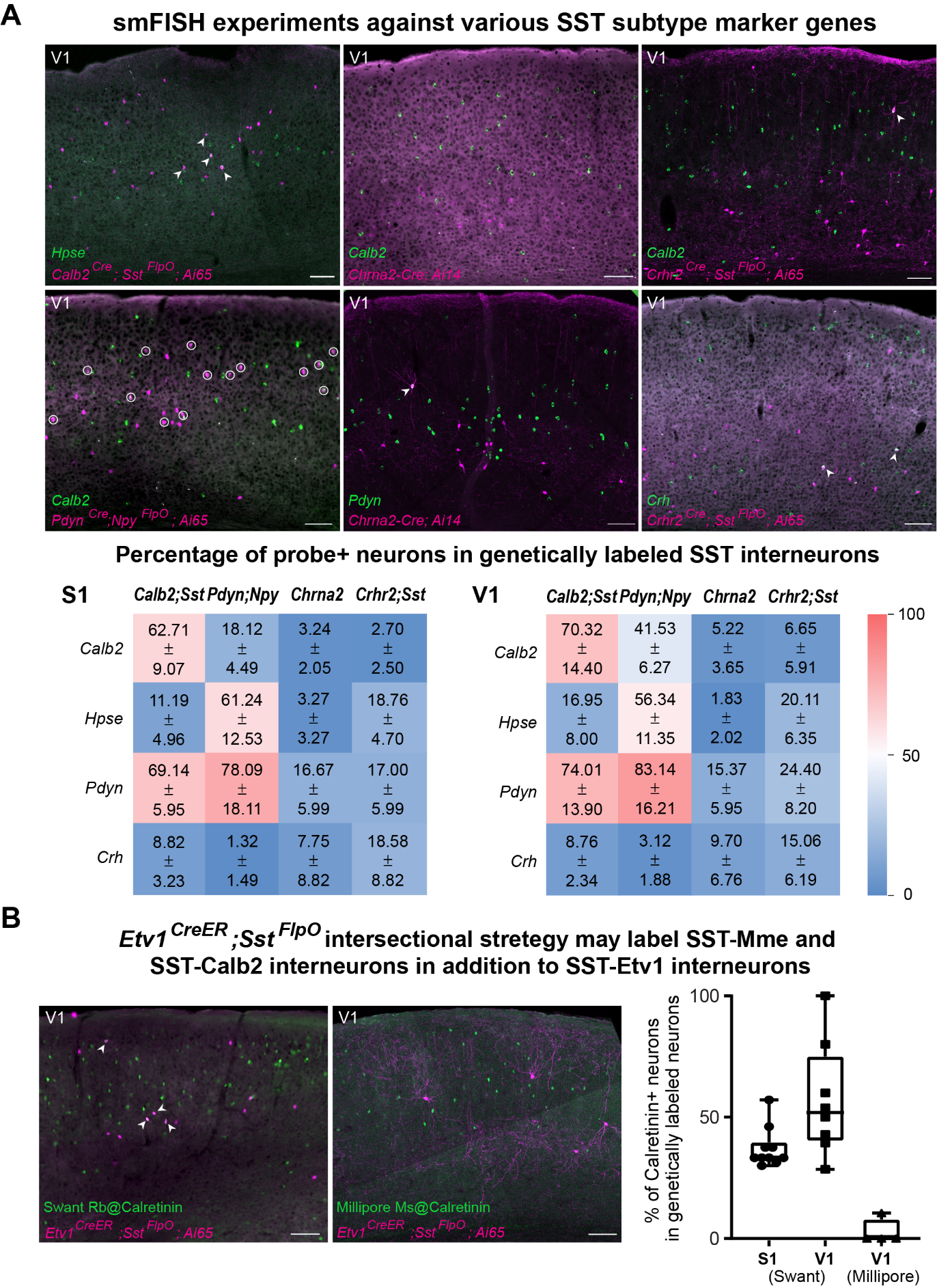

**Figure S5. smFISH experiments against various marker genes for assessing the specificity of genetic targeting strategies. Related to Figure 2.**

(A) Representative images of smFISH experiments against various marker genes on genetically labeled SST interneurons in V1. Circles or arrows indicating genetically labeled neurons (endogenous fluorescence, magenta) that are positive for the probe (green) against different genes. Scale bars, 100 µm. Heatmap showing the quantification of the percentage (mean ± SD) of genetically labelled SST interneurons that showed probe expression in S1 and V1.

(B) Immunostaining against calretinin with two different antibodies showed different degrees of overlapping with genetically labelled SST interneurons. Arrows indicating genetically labeled neurons that are immunostained by antibody against calretinin. Scale bars, 100 µm. Calretinin antibody from Swant labeled many more neurons than the antibody from Millipore. *Etv1^CreER^; Sst^FlpO^* genetic strategy showed little overlapping using Millipore anti-calretinin antibody but ~50% overlapping using Swant antibody, suggesting that this genetic strategy likely label SST interneurons with a low expression of calretinin. Since SST-Mme interneurons express a low level of calretinin, *Etv1^CreER^; Sst^FlpO^* genetic strategy likely label SST-Mme interneurons besides SST-Etv1 interneurons. Furthermore, because this genetic strategy could label a variable amount of SST interneurons depending on the extent of recombination, it is possible that some SST-Calb2 interneurons are also labeled by this genetic strategy with a high degree of recombination.

All quantifications include at least three experiments that were performed on three different mice.

| **Parameter** | **SST-Calb2 mean ± SD n = 21** | **SST-Myh8 mean ± SD n = 12** | **SST-Nmbr mean ± SD n = 13** | **p Value** |
| --- | --- | --- | --- | --- |
| **Vrest (mV)** | ­53.26 ± 0.84 | ­47.27 ± 1.28 | ­57.65 ± 0.95 | **<.0001** |
| **IR (MΩ)** | 311.09 ± 12.25 | 323.64 ± 23.39 | 256.73 ± 24.63 | .097 |
| **Sag ratio** | 0.92 ± 0.01 | 0.91 ± 0.01 | 0.93 ± 0.01 | .128 |
| **AP Amplitude (mV)** | 110.46 ± 1.98 | 97.63 ± 2.7 | 107.14 ± 3.78 | **.007** |
| **AP Half-Width (ms)** | 1.07 ± 0.05 | 1.32 ± 0.1 | 1.02 ± 0.08 | **.029** |
| **AP Max Rise (mV/ms)** | 316.39 ± 14.6 | 230.20 ± 23.52 | 290.08 ± 29.98 | **.025** |
| **AHP Amplitude (mV)** | 14.68 ± 1.23 | 12.69 ± 1.36 | 16.83 ± 1.48 | .177 |
| **AP Threshold (mV)** | ­40.40 ± 0.49 | ­38.99 ± 1.26 | ­38.93 ± 1.03 | .213 |
| **HFF (hZ)** | 52 ± 3.87 | 48.58 ± 6.3 | 74 ± 4.64 | **.004** |
| **Adaptation** | 2.20 ± 0.14 | 2.26 ± 0.12 | 1.69 ± 0.14 | **.042** |
| **Rebound APs** | 1.48 ± 0.46 | 6.42 ± 0.61 | 0 ± 0 | **<.0001** |

**Table S5. Electrophysiological properties of SST-Calb2, SST-Myh8, and SST-Nmbr interneurons in S1. Related to Figure S3.**

Mean ± SEM is reported for all electrophysiology parameters measured. The p value of the F statistic reports which parameters showed significantly different distributions by one‐way ANOVA. Individual mean comparisons with Tukey correction: Vrest: SST-Calb2 vs. SST-Myh8 p = .0002, SST-Calb2 vs. SST-Nmbr p = .01, SST-Myh8 vs. SST-Nmbr p < .0001; AP Amplitude: SST-Calb2 vs. SST-Myh8 p = .005; AP halfwidth: SST-Myh8 vs. SST-Nmbr p = .04; AP Max Rise: SST-Calb2 vs. SST-Myh8 p = .02; HFF: SST-Calb2 vs. SST-Nmbr p = .02, SST-Myh8 vs. SST-Nmbr p = .006; Adaptation: SST-Myh8 vs. SST-Nmbr p = .03; Rebound APs: SST-Calb2 vs. SST-Nmbr p < .0001, SST-Calb2 vs. SST-Nmbr p = .02, SST-Myh8 vs. SST-Nmbr p < .0001.

| **Parameter** | **L2/3 SST-Calb2 mean ± SD n = 10** | **L5a SST-Calb2 mean ± SD n = 11** | **p Value** |
| --- | --- | --- | --- |
| **Vrest (mV)** | ­52.26 ± 1.03 | ­54.17 ± 1.28 | .266 |
| **IR (MΩ)** | 329.43 ± 22.67 | 294.42 ± 30.66 | .377 |
| **Sag ratio** | 0.91 ± 0.01 | 0.92 ± 0.01 | .274 |
| **AP Amplitude (mV)** | 110.39 ± 2.78 | 110.53 ± 2.94 | .973 |
| **AP Half-Width (ms)** | 1.09 ± 0.08 | 1.05 ± 0.05 | .720 |
| **AP Max Rise (mV/ms)** | 316.28 ± 23.94 | 316.49 ± 18.57 | .994 |
| **AHP Amplitude (mV)** | ­15.81 ± 1.74 | ­13.65 ± 1.76 | .395 |
| **AP Threshold (mV)** | ­40.10 ± 0.58 | ­40.68 ± 0.8 | .570 |
| **HFF (hZ)** | 51.9 ± 7.42 | 52.09 ± 3.5 | .981 |
| **Adaptation** | 1.88 ± 0.14 | 2.46 ± 0.19 | **.037** |
| **Rebound APs** | 1.7 ± 0.63 | 1.27 ± 0.68 | .652 |

**Table S6. Electrophysiological properties of SST-Calb2 interneurons in L2/3 and L5 in S1. Related to Figure S3.**

Mean ± SEM is reported for all electrophysiology parameters measured. The p‐value reports which parameters were significantly different across layers by t‐test.

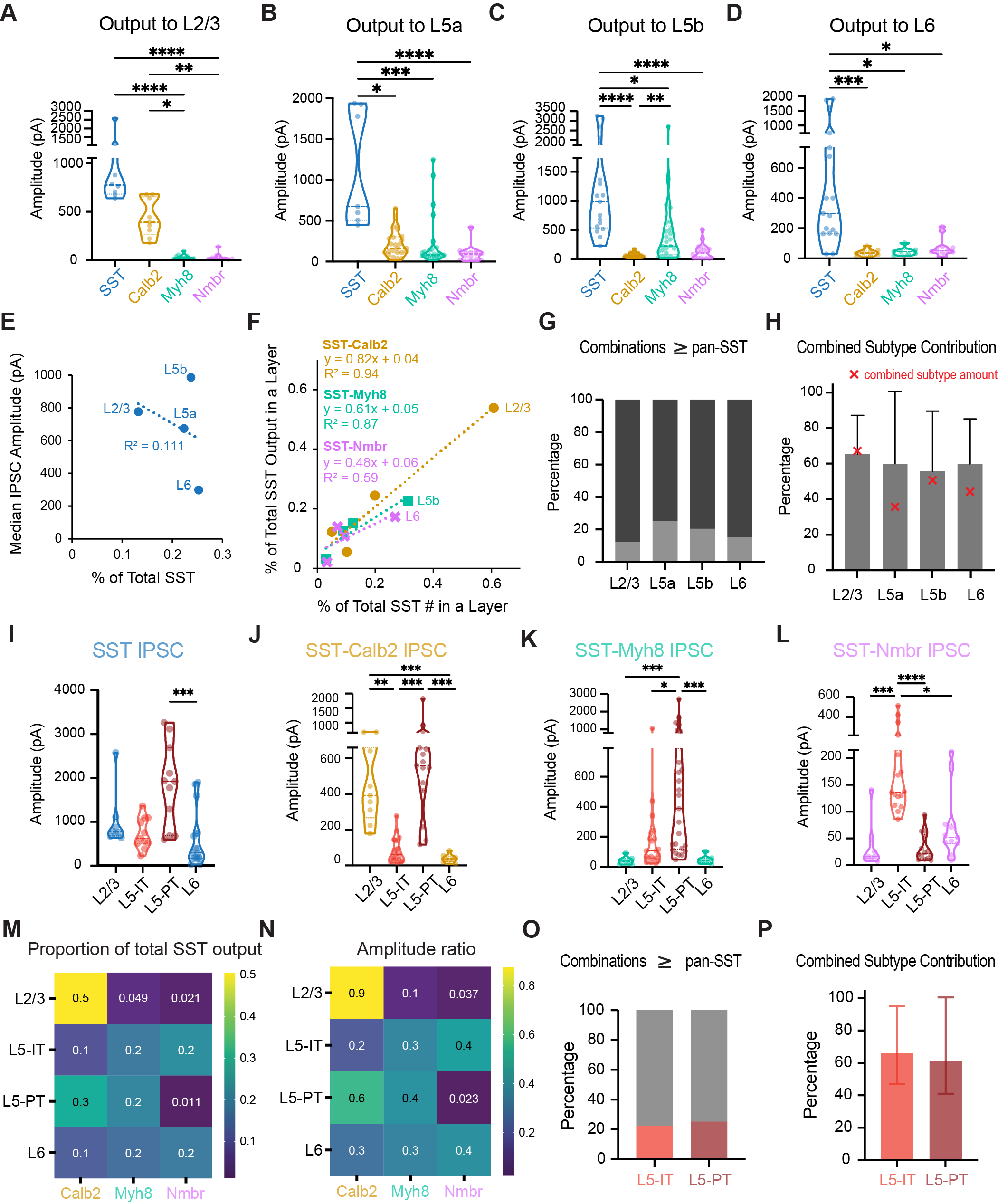

**Figure S6. Pyramidal neurons receive laminar and cell-type specific input from different SST subtypes. Related to Figure 3 and Figure 4.**

(A) Violin plot of evoked IPSC in L2/3 pyramidal neurons. Pan‐SST interneuron response was not significantly greater than SST-Calb2 (p > .999) but was greater than SST‐Myh8 and SST‐Nmbr (p < .0001). SST‐Calb2 was significantly greater than SST‐Myh8 (p = .01) and SST‐Nmbr (p = .009). Kruskal‐Wallis test with Dunn’s correction.

(B) Evoked IPSC in L5a pyramidal neurons. Pan‐SST interneuron response was greater than all three subtypes (pan-SST vs. SST‐Calb2 p = .02, vs. SST‐Myh8 p = .0006, vs. SST‐Nmbr p < .0001). Kruskal‐Wallis test with Dunn’s correction.

(C) Evoked IPSC in L5b pyramidal neurons. Pan‐SST interneuron response was greater than all three subtypes (pan-SST vs. SST‐Calb2 p < .001, vs. SST‐Myh8 p = .02, vs. SST‐Nmbr p < .0001). SST‐Myh8 response was significantly greater than SST‐Calb2 (p = .002) but not SST‐Nmbr (p = .06). Kruskal‐Wallis test with Dunn’s correction.

(D) Evoked IPSC in L6 pyramidal neurons. Pan‐SST interneuron response was greater than all three subtypes (pan-SST vs. SST‐Calb2 p = .0009, vs. SST‐Myh8 p = .01, vs. SST-Nmbr p = .02). Kruskal‐Wallis test with Dunn’s correction.

(E) Percentage of the number of SST interneuron residing in a particular layer out of total number of SST interneurons does not correlate with the percentage of the inhibitory output by total SST interneurons in each layer.

(F) Same plot as Figure 3I with data points from different SST subtypes separately labeled.

(G) Random combinations of SST‐Calb2, SST‐Myh8, and SST‐Nmbr IPSC amplitudes were combined and compared to a pan‐SST evoked IPSC amplitude (see Methods). Graph of the percent of simulations where the difference was below (light grey) and above (dark grey) zero. The combined evoked IPSC amplitude from SST-Calb2, SST-Myh8, and SST-Nmbr was smaller than the pan-SST response in L2/3 on 87.68% of the trials, in L5a on 74.85% of trials, in L5b on 79.58% of the trials, and in L6 on 84.86% of the trials.

(H) The proportion of the pan‐SST response accounted for by a linear combination of SST‐Calb2, SST‐Myh8, and SST‐Nmbr inputs. Graph of median ratios. Error bars are interquartile range. Red crosses indicate the percentage of the combined amount of three SST subtypes found in each layer out of total SST interneurons.

(I) Quantification of the evoked IPSC amplitude upon pan-SST stimulation across pyramidal neuron types. L5-PT IPSC was significantly greater than L6 (p = .0005), all other comparisons not significant (L2/3 vs. L6 p = .11520, L5-IT vs. L5-PT p = .0870, rest p > .9999). Kruskill-Wallis test with Dunn’s correction.

(J) As in (I) for SST-Calb2 interneurons. IPSC was significantly greater in L2/3 and L5-PT neurons than L5-IT or L6 neurons (L2/3 vs. L5-IT p = .0019, L2/3 vs. L6 p = .0008, L5-PT vs. L5-IT and L5-PT vs. L5 p < .0001. L2/3 vs. L5-PT not significant, p > .9999). Kruskill-Wallis test with Dunn’s correction.

(K) As in (I) for SST-Myh8 interneurons. IPSC in L5-PT neurons was significantly greater than in all other cell types (L2/3 vs. L5-PT and L5-PT vs. L6, p < .0001. L5-IT vs. L5-PT p = .0190). Kruskill-Wallis test with Dunn’s correction.

(L) As in (I) but for SST-Nmbr interneurons. IPSC in L5-IT neurons was significantly greater than in all other cell types (L2/3 vs. L5-IT p = .0002, L5-IT vs. L5-PT p < . 0001, L5-IT vs. L6 p = .0424). Kruskill-Wallis test with Dunn’s correction.

(M) Heatmap of the proportion of inhibition from individual SST subtype as compared to the inhibition from pan-SST interneurons in different layers and pyramidal neuron cell types.

(N) The ratio of IPCS amplitude for the three SST subtypes with the combined median IPSC amplitude of the three subtypes normalized to 1 for particular layer or pyramidal neuron cell type.

(O) Random combinations of SST‐Calb2, SST‐Myh8, and SST‐Nmbr IPSC amplitudes were combined and compared to a pan‐SST evoked IPSC amplitude (see Methods). Graph of the percent of simulations where the difference was below (pink/red) and above (grey) zero.

(P) The proportion of the pan‐SST response accounted for by a linear combination of SST‐Calb2, SST‐Myh8, and SST‐Nmbr inputs. Graph of median ratios. Error bars are interquartile range.

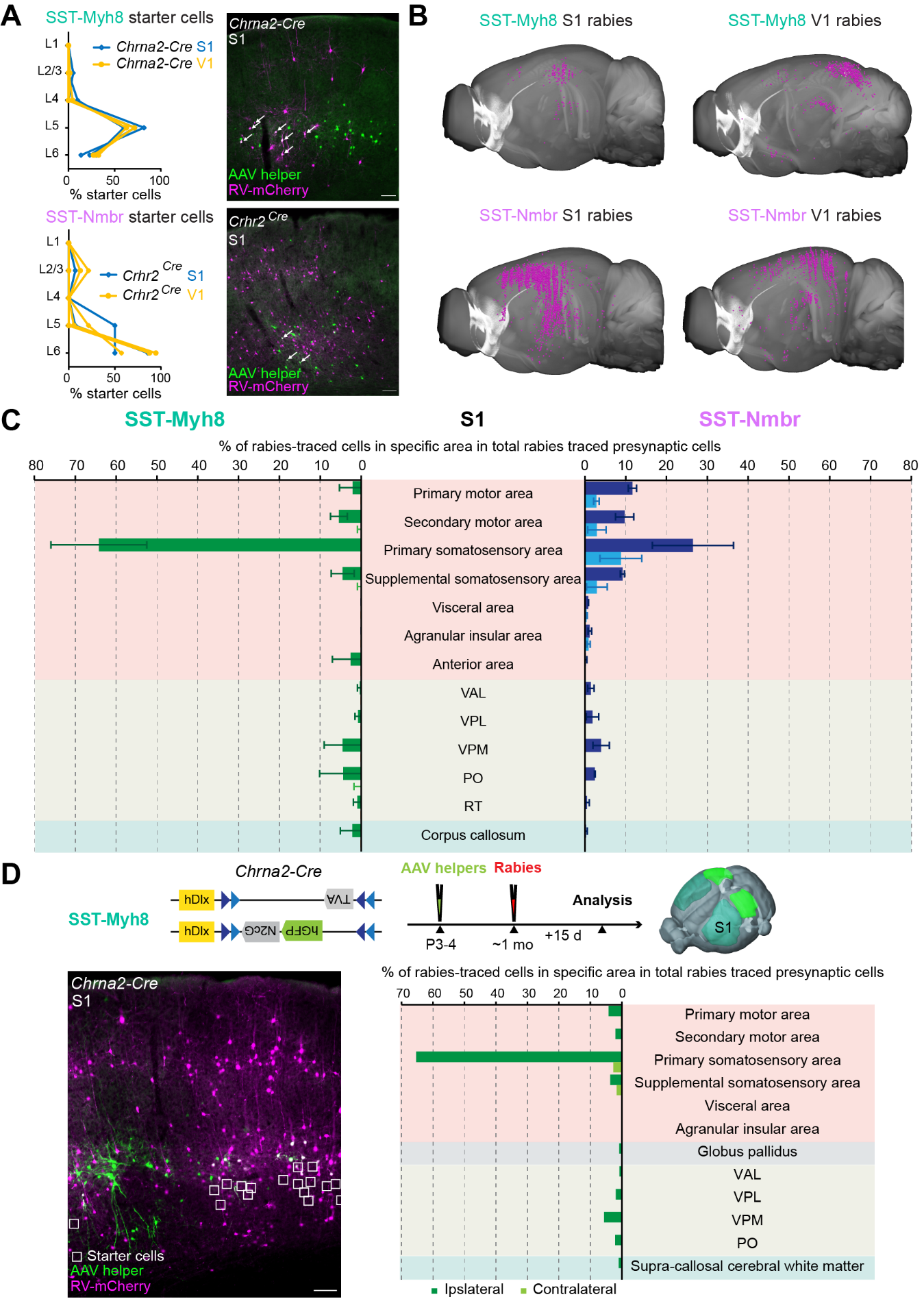

**Figure S7. Monosynaptic rabies tracing method, rabies tracing results in S1, and one test experiment using the matching AAV helper viruses for rabies tracing from SST-Myh8 subtype as used for SST-Nmbr. Related to Figure 6.**

(A) (left) Laminar distribution of rabies-infected starter cells for rabies tracing experiments from SST-Myh8 and SST-Nmbr interneurons in S1 and V1. (right) Representative images showing starter cells of SST-Myh8 and SST-Nmbr interneurons. Neurons expressing AAV-helpers are in green, Rabies (RV) in magenta and starter cells (arrows) are identified by both channels. Scale bar, 100 µm.

(B) Representative examples of rabies retrograde labeling from SST-Myh8 and SST-Nmbr interneurons in S1 and V1 using Neuroinfo 3D rendering. Magenta dots represents the location of rabies traced presynaptic neurons.

(C) Presynaptic inputs to SST-Myh8 and SST-Nmbr interneurons in S1 quantified as the percentage of rabies traced cells in each regional category out of the total number of cells labeled in the brain (n = 3 for SST-Myh8, n = 2 for SST-Nmbr). Top 10 input regional categories for either SST subtype are included in the plot.

(D) A rabies tracing experiment of SST-Myh8 interneurons using the same AAV-helper viruses used for rabies tracing experiments for SST-Nmbr interneurons. (top) Experimental design of the experiment. The design of AAV-DIO-helper viruses and the timeline of AAV-helpers and N2cRV injections for tracing from SST-Myh8 in S1 using *Chrna2-Cre* mouse line is illustrated. Rabies tracing patterns were analyzed 15 days post-infection. (bottom left) Representative image showing the starter cells. (bottom right) Presynaptic inputs identified were quantified as the percentage of rabies traced cells in each regional category out of the total number of cells labeled in the brain. Top 10 input regional categories are included in the plot.

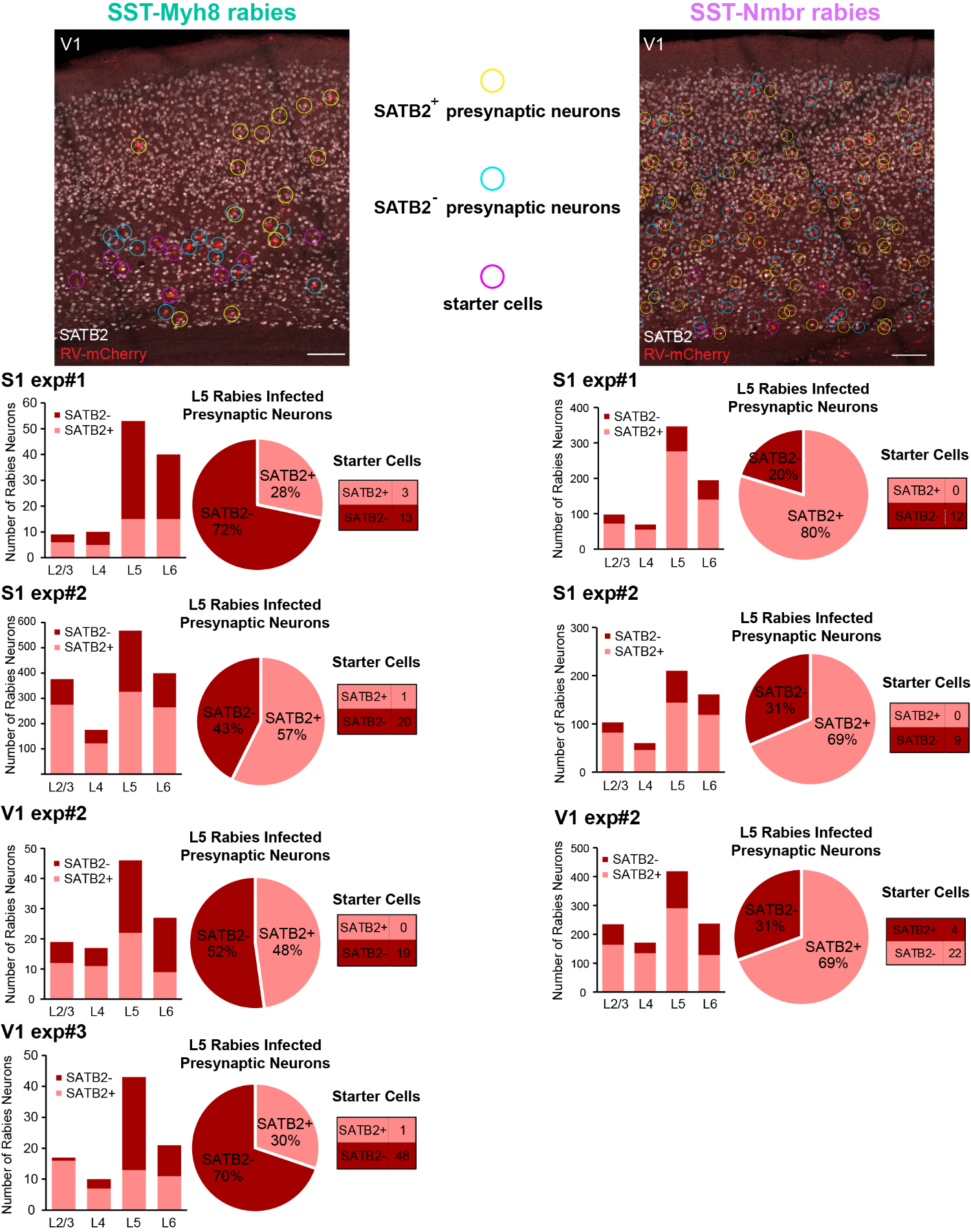

**Figure S8. Identity of the rabies traced local presynaptic neurons to two SST subtypes. Related to Figure 6.**

(left column from top to bottom) Representative image of immunostaining against SATB2 (white) on a brain slice with rabies traced presynaptic neurons (red) targeting SST-Myh8 interneurons in V1. Yellow circles indicate SATB2+ rabies traced presynaptic neurons. Cyan circles indicate SATB2- rabies traced presynaptic neurons. Magenta circles indicate starter cells. Below this are quantification from individual rabies tracing examples. For each experiment, (left) the histogram shows the number of rabies traced neurons in each layer, (middle) the pie chart shows the percentage of SATB2+ and SATB2- neurons labeled by rabies in L5, and (right) the table shows the number of starter cells. As expected, most of the starter cells are Satb2- interneurons, though occasionally there is small contamination of a low number of SATB2+ starter cells.

(right column) same elements for rabies tracing experiments from SST-Nmbr interneurons.
